# The role of mechanical interactions in EMT

**DOI:** 10.1101/2020.12.09.418434

**Authors:** Ryan J. Murphy, Pascal R. Buenzli, Tamara A. Tambyah, Erik W. Thompson, Honor J. Hugo, Ruth E. Baker, Matthew J. Simpson

## Abstract

The detachment of cells from the boundary of an epithelial tissue and the subsequent invasion of these cells into surrounding tissues is important for cancer development and wound healing, and is strongly associated with the epithelial-mesenchymal transition (EMT). Chemical signals, such as TGF-*β*, produced by surrounding tissue can be up-taken by cells and induce EMT. In this work, we present a novel cell-based discrete mathematical model of mechanical cellular relaxation, cell proliferation, and cell detachment driven by chemically-dependent EMT in an epithelial tissue. A continuum description of the model is then derived in the form of a novel nonlinear free boundary problem. Using the discrete and continuum models we explore how the coupling of chemical transport and mechanical interactions influences EMT, and postulate how this could be used to help control EMT in pathological situations.

## 1 Introduction

Cell detachment driven by epithelial-mesenychmal transitions (EMT) is crucial to many biological processes: embryonic development; later development in adults; wound healing; and cancer development [1, 2, 3]. During EMT, changes in gene expression and post-translational regulation mechanisms lead to increased invasive ability through the loss of epithelial characteristics and the acquisition of mesenchymal characteristics [1]. This transition is characterised by the loosening of cell-cell junctions and breakdown of the basement membrane [1]. EMT can be induced by chemical signals, such as TGF-*β* [4], and is regulated by physical signals such as mechanical stress [5]. While EMT plays an important role in development, where it is a highly controlled and regulated process, EMT can be detrimental when initiated by cancer systems as it accelerates malignant progression and metastasis [6]. Furthermore, as 90% of cancer related deaths are associated to metastatic spread rather than cancer limited to a primary site [7], EMT is an important factor when considering therapy regimes [8, 9, 10, 11]. Previous theoretical models of EMT have largely neglected the role of mechanical interactions. Therefore, in this work we develop a novel mathematical model to explore how mechanical interactions between cells influence EMT and the evolution of a primary tumour. Using our model, we ask when do tumours grow or shrink, and how mechanical interactions and an EMT-inducing chemical could influence the rate of cell detachment. These insights could be used to help understand how to control EMT in pathological situations.

Mathematical models have proven to be a powerful tool to improve our understanding of EMT by providing a conceptual framework in which to integrate and analyse experimental data and make testable predictions, some of which have since been experimentally validated, for example, the existence of the epithelial/mesenchymal hybrid state [3, 4, 12, 13, 14, 15] and waves of temporal cell-cell detachments [16, 17]. Experimental and modelling studies are typically performed either at the single-cell level, by considering regulatory networks [3, 12, 13], or at the population-level, for example where cell populations are modelled with lattice based frameworks and the inclusion of cell-cell communication results in spatial heterogeneity [18]. However, existing models for EMT typically do not account for mechanical relaxation nor cell proliferation, both of which influence cell migration and cell size [5, 18]. These processes are thought to play a key role in cell-cell communication, tissue size, and the rate of cell detachment driven by EMT [18, 19]. Further, existing models typically do not connect descriptions of single-cell processes to population-scale behaviours.

In this work, we develop and explore a novel mathematical model of EMT which includes mechanical cellular relaxation, cell proliferation and cell detachment driven by chemical signals. We allow individual cells to detach from the tissue at the free boundary where the chemical concentration is highest [18]. This leads to a novel nonlinear free boundary problem. The evolution of epithelial monolayers and tumours have previously been modelled as free boundary problems [20, 21, 22]. However, many previous studies specify a classical one-phase Stefan condition [23] at the free boundary, where the rate of expansion of the free boundary is assumed proportional to the spatial gradient of the density without strong biological justification, or the evolution of the tissue length is specified according to experimental observations [24, 25, 26, 27, 28, 29]. Here, the evolution of the tissue boundary arises naturally from the cell-scale processes of cell proliferation, mechanical cellular relaxation [30, 31, 32], and cell detachment.

To implement this model, we start with a cell-based discrete model, where we prescribe individual cell-level properties, and then derive the corresponding tissue-level continuum partial differential equation model. This approach extends previous studies [30, 31, 32, 33,34, 35, 36, 37, 38] all of which consider mechanical cell movement, but do not consider cell detachment driven by EMT. The continuum model is useful to analyse possible behaviours of the model including tissue shrinkage, tissue homeostasis, and tissue growth depending on the initial number of cells, mechanical cell properties, the rate of proliferation, and chemical diffusivity. Importantly, we provide guidance when the discrete and continuum models are accurate. To simulate the mathematical model numerically, we devise a new method to incorporate chemical diffusion in an evolving population of cells with variable cells lengths. This work is structured in the following way; we present the cell-based discrete model (Section 2.1) and derive the corresponding continuum model (Section 2.2). Then using the new discrete and continuum models we explore when tumours grow or shrink, and how mechanical interactions and an EMT-inducing chemical could influence the rate of cell detachment. To do so we consider different mechanisms in the models: cell-length-independent proliferation mechanism and chemically-independent cell detachment (Section 3.1); cell-length-dependent proliferation mechanism and chemically-independent cell detachment (Section 3.2); chemically-dependent cell detachment driven by an EMT-inducing chemical which diffuses slowly (Section 3.3), or which diffuses quickly (Section 3.4).

## 2 Model description

In this section, we present the new discrete model of mechanical cellular relaxation, cell proliferation, and cell detachment driven by a chemically-dependent EMT process in an epithelial tissue. We then derive the corresponding continuum model. To simplify the model, we suppose that linear diffusion is the key transport mechanism by which the chemical that induces EMT is transported from the free boundary inwards through the cells in the epithelial tissue (Figure 1).

**Figure 1.**
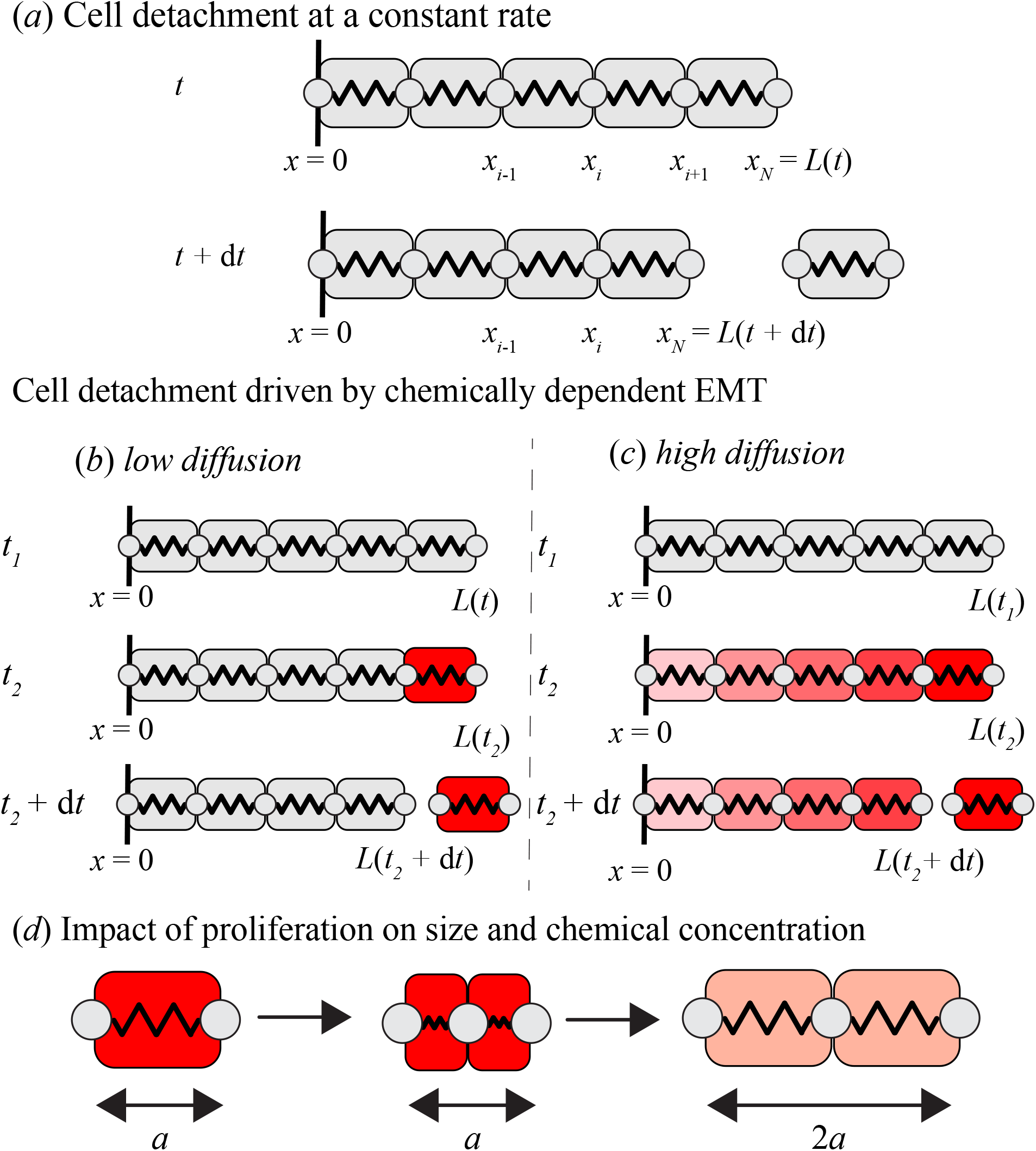
Schematic for models of EMT and cell detachment. (a) Boundary cell detachment at a constant rate. (b)-(c) Boundary cell detachment driven by chemically-dependent EMT with (b) low diffusion and (c) high diffusion. The EMT-inducing chemical diffuses inwards from the external environment through the cell at the free boundary. In (b) with low diffusion the boundary cell contains most of the EMT-inducing chemical whereas in (c) with high diffusion the chemical is spread throughout the boundary and internal cells. Chemical concentration is shown with colouring: low concentration below the cell-detachment chemical threshold (light red) to a higher concentration above the cell-detachment chemical threshold (dark red). (d) Proliferation produces two identical daughter cells. Each daughter cell mechanically relaxes to the resting cell length and the concentration changes accordingly.

### 2.1 Discrete model

We consider a one-dimensional chain of cells to represent the cross-section of an epithelial tissue (Figure 1). Each cell is assumed to act like a mechanical spring [22, 30, 31, 32, 35, 36]. The tissue has a fixed boundary at *x*= 0 and a free boundary at *x* = *L*(*t*) *>* 0. Cells undergo mechanical relaxation which results in changes in cell length and corresponding movements of cell boundaries. Cell *i*, for *i* = 1, 2, …, *N* (*t*), occupies the interval *x*_*i*_(*t*) *< x < x*_*i*+1_(*t*), where *x*_*i*_(*t*) and *x*_*i*+1_(*t*) are the positions of the left and right boundaries of the cell, respectively, so that cell *i* has length *l*_*i*_(*t*) = *x*_*i*+1_(*t*) *− x*_*i*_(*t*). Each cell is prescribed with cell stiffness *k >* 0 and resting cell length *a >* 0. We assume that the motion of each cell boundary is subject to mechanical interactions and occurs in a viscous medium, resulting in a drag force with drag coefficient *η >* 0 [22, 35, 39, 40]. Then, as cells move in dissipative environments, the motion is assumed to be overdamped [35, 39] and the cell boundary evolution equations are

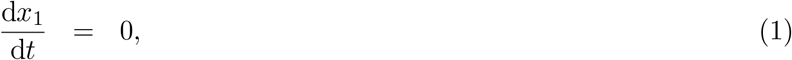

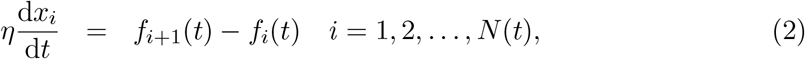

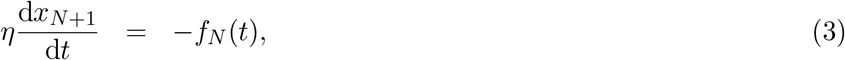

where, for simplicity, we use *f*_*i*_(*t*) = *k* (*l*_*i*_(*t*) − *a*) to represent a linear Hookean force law, and *N* (*t*) evolves in time due to proliferation and cell detachment.

We assume any cell in the tissue is able to proliferate via a stochastic cell proliferation mechanism that may depend on cell length. We assume that cell *i* proliferates with probability *P* (*l*_*i*_)d*t* in the small time interval [*t, t* + d*t*), where the function *P* (·) depends on the proliferation mechanism considered, and *l*_*i*_ is the current cell length (Figure 2(b)). When a proliferation event occurs in a cell of length *l*_*i*_, the cell divides into two cells of length *l*_*i*_*/*2, and any chemical inside the cell is conserved and divided equally between the two daughter cells. The chemical concentration subsequently dilutes as the daughter cells mechanically relax to their resting cell lengths (Figure 1(d)).

**Figure 2.**
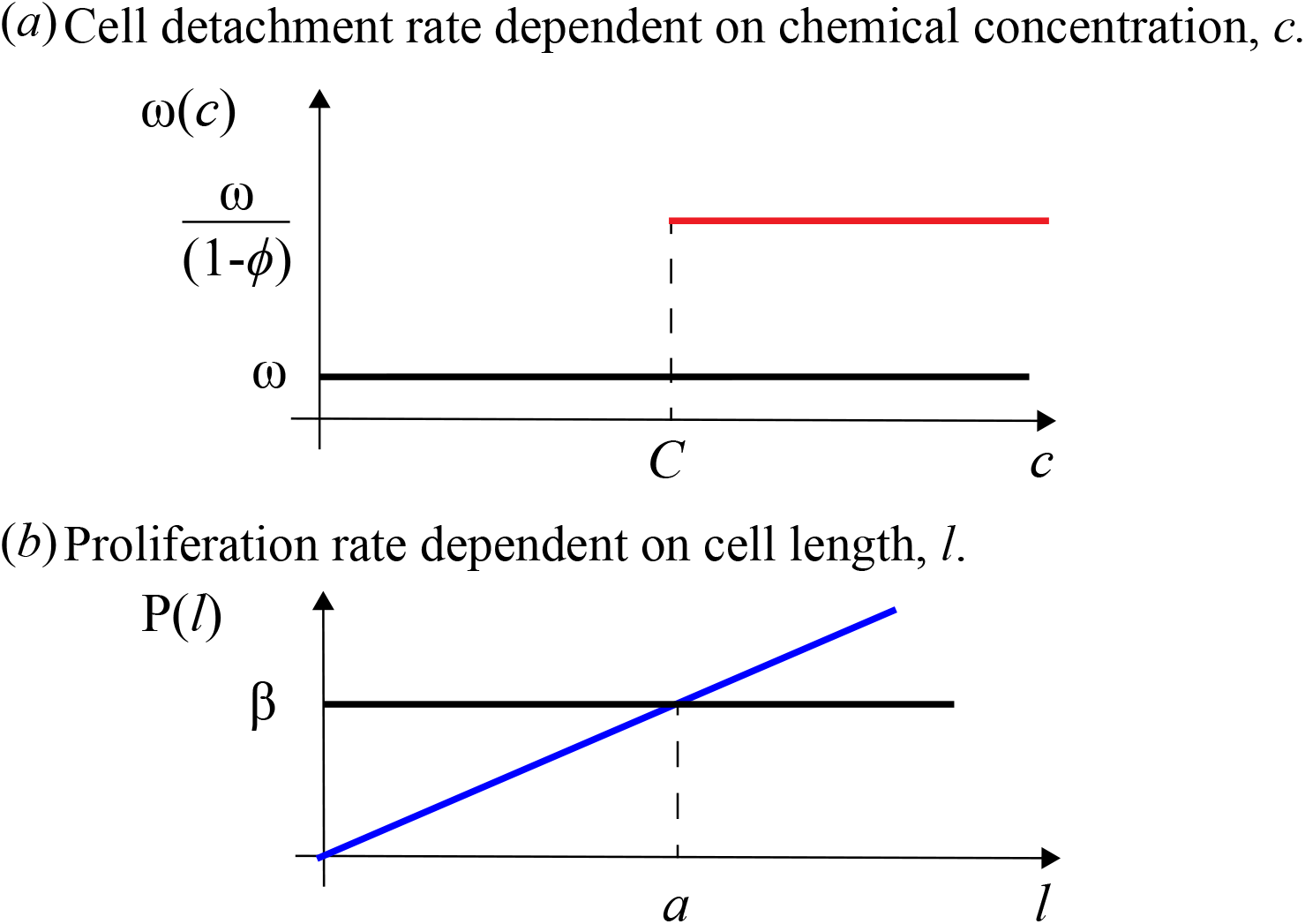
Cell detachment driven by chemically-dependent EMT and proliferation mechanisms. (a) Cell detachment mechanisms: cell detachment at a constant rate *ω* Independent of chemical concentration (black); cell detachment at a constant rate *ω/*(1 − *ϕ*) when the concentration, *c*, is above a concentration threshold, *C* (red). (b) Cell-length-dependent proliferation mechanisms: independent of cell length at rate *β* (black); linearly dependent on cell-length (blue) defined by ensuring *P* (0) = 0 and *P* (*a*) = *β*. In this main manuscript we set *ω* = 0.1, *ϕ* = 0.9, *C* = 500 and *β* = 0.0025, and vary *ϕ* in Appendix D.6.

We assume an external source of EMT-inducing chemical and suppose that linear diffusion is the key transport mechanism by which the chemical is transported into and through the epithelial tissue. To model linear diffusion we consider the chain of cells to be a non-uniform lattice on which we can simulate a point-jump process for molecules of the chemical. As it is computationally expensive to track many individual particles, we focus on the chemical concentration. In the epithelial tissue, each cell *i* has a chemical concentration *c*_*i*_(*t*) = *M*_*i*_(*t*)*/l*_*i*_(*t*), where *M*_*i*_(*t*) is the number of molecules of the chemical in cell *i* at time *t*, and *l*_*i*_(*t*) is the length of cell *i* at time *t*. Therefore, the concentration in each cell is assumed to be well-mixed. Then the equations governing chemical concentration are [41]

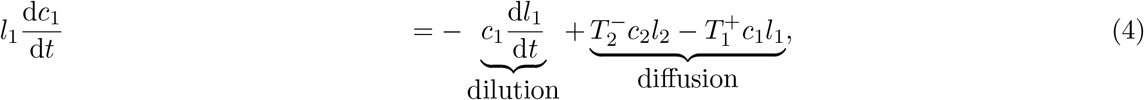

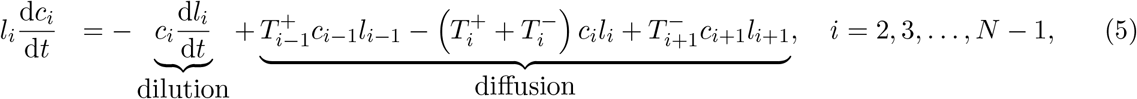

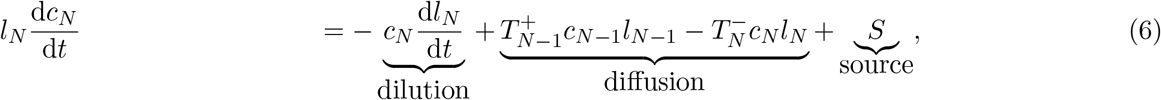

where 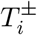 are left and right transition rates that model linear diffusion of chemical molecules between neighbouring cells with diffusivity *D*, respectively. How these transition rates are chosen requires great care and is detailed in Appendix B where we introduce a new method, called the *Interval-Voronoi* method. The dilution term in Equations (4)-(6) represents the fact that chemical concentrations increase/decrease as the cell length reduces/increases. To mimic a chemical diffusing into the tissue from an external source, we assume that a constant number of molecules per unit time, *S*, is provided to cell *N* from the external source. We further assume that the chemical cannot diffuse across the boundary at the rear of the tissue (*x* = 0). This assumption corresponds to modelling only the right hand side of an epithelial tissue where *x* = 0 is the middle of the tissue.

Given the chemical concentrations in each cell, we introduce cell detachment driven by EMT which is a key feature of the discrete mechanical model. Here, we consider cell detachment to be a two-step process. The first step models the EMT process itself as a cell-state transition whereby cells acquire an invasive phenotype. The boundary cell gains the invasive phenotype when its chemical concentration is above a constant threshold, *C* (Figure 2)[4, 12, 13]. If at any time the chemical concentration drops below the threshold the cell loses its invasive phenotype, which it can only regain once the chemical concentration increases above *C*. The ability of the cell to gain and lose its invasive phenotype is associated with epithelial-mesenchymal plasticity [14]. The second part of the process is where the boundary cell, once it acquires the invasive phenotype, detaches from the tissue [42] in [*t, t* + d*t*) with probability *ω*(*c*_*N*_ (*t*)) d*t* where *c*_*N*_ (*t*) is the chemical concentration in the boundary cell *N* at time *t*. Once a cell detaches we no longer consider its dynamics and we assume it moves away from the epithelial tissue.

As we are interested in whether tumours grow or shrink, we can consider the evolution of the total number of cells, *N* (*t*), which depends on the balance between proliferation and EMT. For an individual realisation of this discrete model, *N* (*t*) is expected to increase when

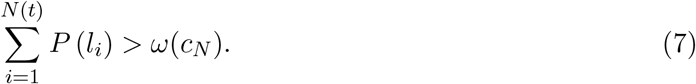

We numerically simulate the discrete model governed by Equations (1)–(6), and prescribe initial conditions for the cell positions, *x*_*i*_(0), the mechanical cell properties *k* and *a*, drag coefficient *η*, as well as proliferation properties, and assume that there is initially no chemical inside any cell in the tissue (Appendix B).

### 2.2 Continuum model

We now derive the corresponding free boundary continuum model for cell detachment driven by chemically-dependent EMT. Components of this model have been derived in our previous studies, and where this is the case we state the equation and provide a reference to the reader for full details [30, 32, 35, 36].

The cell density, *q*(*x, t*) *>* 0, which is the number of cells per unit length and the continuous analogue of 1*/l*_*i*_, evolves according to the following nonlinear moving boundary problem [36]

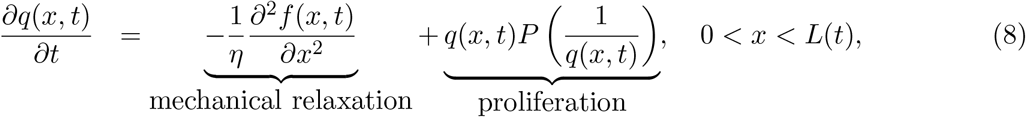

where *f* (*x, t*) = *k* (1*/q*(*x, t*) *− a*) is the continuous analogue of the discrete Hookean force law and *P* (1*/q*(*x, t*)) is the continuous analogue of *P* (*l*_*i*_).

The fixed boundary condition at *x* = 0, corresponding to Equation (1), is [35]

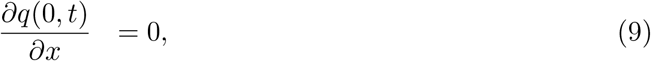

and the mechanical relaxation condition at the free boundary, *x* = *L*(*t*), gives rise to a nonlinear boundary condition [30, 32]

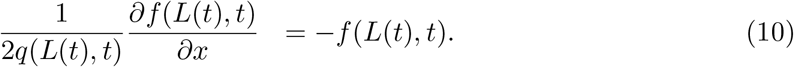

The boundary conditions in Equations (9) and (10) ensure that no cells are lost by crossing the tissue boundaries but cells can still detach at the free boundary, *x* = *L*(*t*). In the continuum model this corresponds to loss of tissue material at a moving interface [43]. To capture cells detaching from the tissue at *x* = *L*(*t*), we consider conservation of mass and derive the following evolution equation for the free boundary (Appendix A)

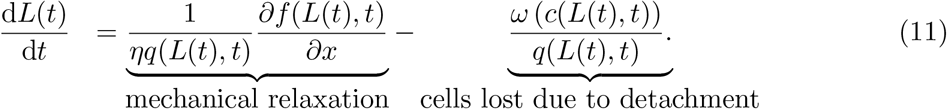

The chemical concentration is governed by the following advection-diffusion equation [24, 25, 32, 44]

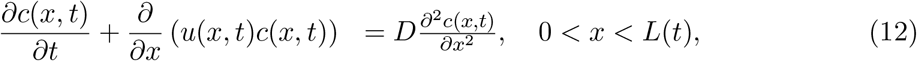

where *D* is the diffusion coefficient, and the cell velocity, *u*(*x, t*), determined from Equation (8), is

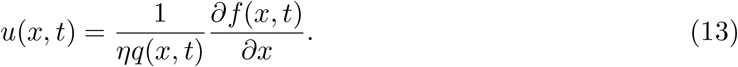

The boundary at *x* = 0 is fixed, and because there is no chemical transport across this boundary in the discrete model, we impose the following no-flux boundary condition

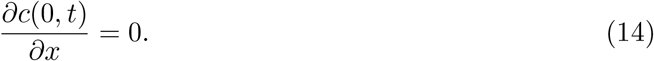

At the free boundary, *x* = *L*(*t*), the only transport of chemical in the discrete model is the supply of a constant number of molecules per unit time, *S*, from the external environment.

The corresponding boundary condition at *x* = *L*(*t*) in the continuum model is

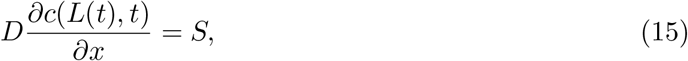

obtained by enforcing that the total flux of *c*(*x, t*) in the frame of reference co-moving with *L*(*t*) is equal to *−S* [32].

We supplement the continuum model with initial conditions for tissue length, *L*(0), density *q*(*x*, 0) for 0 < *x* < *L*(0), and chemical concentration *c*(*x*, 0) for 0 < *x* < *L*(0). Then Equations (8)–(10),(11), and (12)–(15) are solved numerically using a boundary fixing transformation [45] and an implicit finite difference approximation, see Appendix C for further details.

## 3 Results and discussion

The evolution of the number of cells in the epithelial tissue, *N* (*t*), the length of the epithelial tissue, *L*(*t*), and the number of cells that undergo EMT and detach from the epithelial tissue are coupled. Biologically this coupling is of great interest. In the context of cancer, a primary tumour site without EMT is a localised problem, whereas a single tumour site with EMT and cell detachments may result in many secondary tumour sites that can be a greater problem as 90% of cancer related deaths are associated with metastatic spread [7]. We are interested in how mechanical interactions influence EMT and subsequently how tumours grow or shrink. Therefore, we choose parameters to explore the range of behaviours that our new mathematical model of EMT predicts. Further, as the continuum model is useful to analyse possible behaviours, we seek to understand when the continuum model is a good description of the underlying discrete model by considering initial populations with low numbers of cells. Our parameter choices are also consistent with experimental observations that can vary greatly depending on the cell type and driving mechanisms: a cell can proliferate on the order of once every 12 hours to once every few days; EMT can occur over the course of hours, a few days [46], or many days (e.g. 9-12 days [47]); the rate of mechanical relaxation is faster than the rate of proliferation and EMT, with *η/k ≈* 5–16 minutes being a typical experimental value [40]; and a typical experimental value for the resting cell length being *a ≈* 10 µm [33].

### 3.1 Cell-length-independent proliferation and chemically-independent EMT

The simplest model is chemically-independent cell detachment of the boundary cell at a constant rate, *ω*, with cell-length-independent proliferation for each cell at a constant rate, *β* (Figure 2(a),(b)). It is useful to first examine this problem with the continuum model. Conservation of mass (Equation (17)) gives a simple ordinary differential equation for the evolution of *N* (*t*),

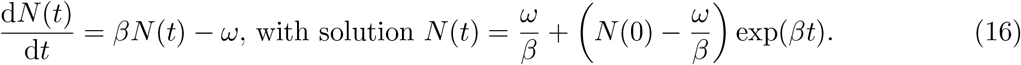

Then, depending on the initial number of cells, *N* (0), and the critical cell number, *ω/β*, there are three possible long-term outcomes: i) *N* (*t*) *→* 0 in finite time, which we refer to as extinction, when *N* (0) *< ω/β* ; ii) *N* (*t*) remains constant for all *t* when *N* (0) = *ω/β*; iii) *N* (*t*) *→ ∞* as *t → ∞* when *N* (0) *> ω/β*. It is clear from Equation (16) that in all cases mechanical interactions do not influence *N* (*t*). However, each cell mechanically relaxes towards its equilibrium length as *N* (*t*) evolves over time. To capture the evolution of the tissue length, *L*(*t*), and cell density, *q*(*x, t*) we solve the full continuum model, governed by Equations (8)-(10) and (11) (Figure 3). The total number of cells which detach grows linearly with time at rate *ω* for chemically-independent EMT when the tissue is not close to extinction, and plateaus if extinction occurs (Figure 10).

**Figure 3.**
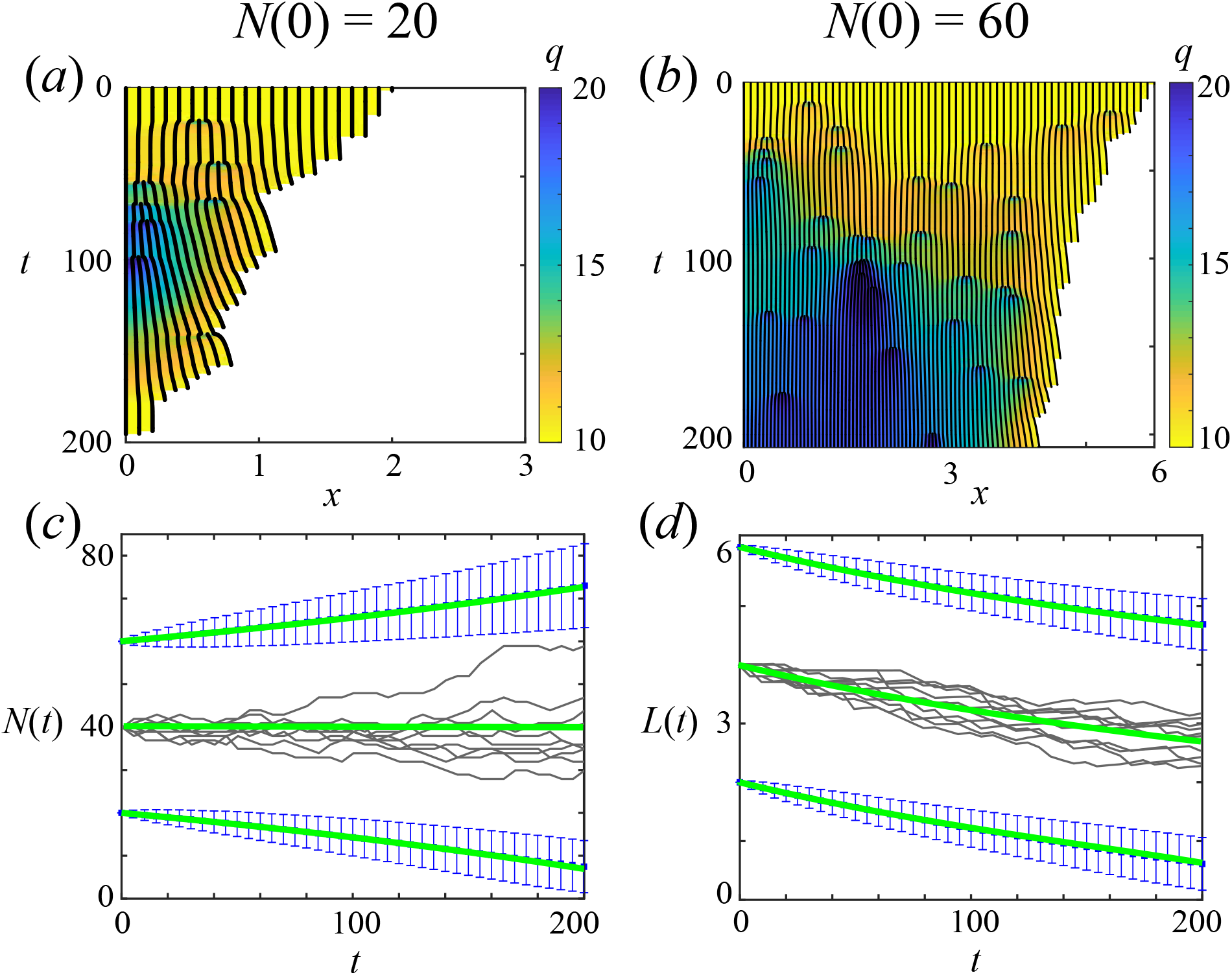
Chemically-independent cell detachment at a constant rate with a cell-length-independent proliferation mechanism. (a)-(b) Kymographs showing evolution of cell boundaries (black curves, note bifurcations when cells divide) for one discrete realisation with cell density, *q*(*x, t*), colouring for (a) *N* (0) = 20 and (b) *N* (0) = 60. Note domain size in (b) is double that of (a). (c)-(d) Three initial cell populations starting at mechanical equilibrium, *L*(0) = *N* (0)*a*. For *N* (0) = 20, 60 the average of 2000 discrete realisations (blue) are compared with the continuum model (green). For *N* (0) = *ω/β* = 40, 10 discrete realisations (grey) are compared with the continuum model (green). (c) Evolution of number of cells, *N* (*t*). (d) Evolution of tissue length, *L*(*t*). Mechanical parameters: *k* = 1, *a* = 0.1, *η* = 1.

In general, the solution of the continuum model provides an accurate approximation for the evolution of *N* (*t*), *L*(*t*) and *q*(*x, t*) when compared to appropriately averaged quantities from many discrete realisations (Figures 3, 11-14). This correspondence between the discrete and continuum model holds provided that the rate of mechanical relaxation, determined by the ratio of cell stiffness to drag coefficient, *k/η*, is sufficiently fast relative to the rate of cell proliferation, see Section 3.3 of [36]; and that *N* (*t*) is sufficiently large to define a continuum, see Appendix D.4. However, when *N* (0) is close to *ω/β* the behaviour of the discrete and the continuum model may differ. Extinction behaviour of the continuum model is deterministic and solely determined by *N* (0), whereas stochastic effects in the discrete model imply that different realisations for the same *N* (0) may sometimes result in extinction and sometimes in unbounded growth (Figures 3(c), Equation (7)) [48]. To quantify this difference between the models, we simulate many identically-prepared realisations of the discrete model and calculate the survival probability of the tissue: the probability that an individual realisation is not extinct at a certain time. By comparing the survival probability of the tissue from the discrete and continuum models for a range of *N* (0), Figure 4 shows that when *N* (0) is close to *ω/β* and when *N* (0) is close to extinction, results from the continuum model may not reflect the behaviour of individual discrete realisations.

**Figure 4.**
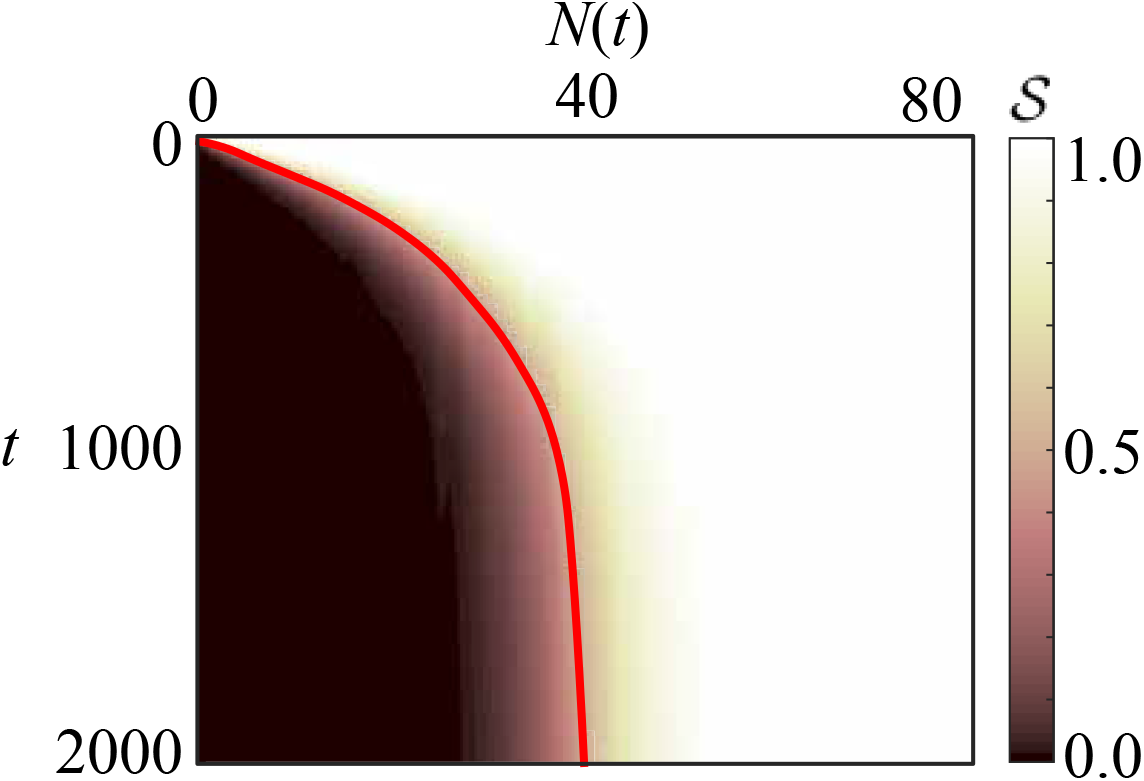
Survival probability of the tissue, *S*, for the cell-length-independent proliferation mechanism and chemically-independent cell detachment with *N* (0) = 1, 2, …, 80. Comparison between the deterministic continuum model (red solid line), and the average of 2000 realisations of the stochastic discrete model (shading). Here, *w/β* = 40.

### 3.2 Cell-length-dependent proliferation and chemically-independent EMT

With a constant cell-length-independent proliferation rate, *N* (0) determines the long-term solution of the continuum model, whereas with cell-length-dependent proliferation we must also consider the initial cell lengths, *l*_*i*_(0), resting cell length, *a*, and the ratio of cell stiffness to drag coefficient, *k/η*. If we consider a linear proliferation mechanism *P* (*l*_*i*_) = *βl*_*i*_*/a*, shown in Figure 2(b), then the critical tissue length is *ωa/β* (Equation (7)). Therefore in this case mechanical interactions between the cells are important. Parameter combinations that lead to extinction with cell-length-independent proliferation may now grow without bound with cell-length-dependent proliferation (Section D.3). Similarly, parameter combinations that lead to unbounded growth with cell-length-independent proliferation may now lead to extinction with cell-length-dependent proliferation. Further, the model predicts that compressed tissues can go extinct faster (Figure 5(a),(c),(e)) than stretched tissues (Figure 5(b),(d),(f)). Good agreement between the continuum model and appropriately averaged quantities from many discrete realisations is also observed when considering cell-length-dependent proliferation (Appendix D.7).

**Figure 5.**
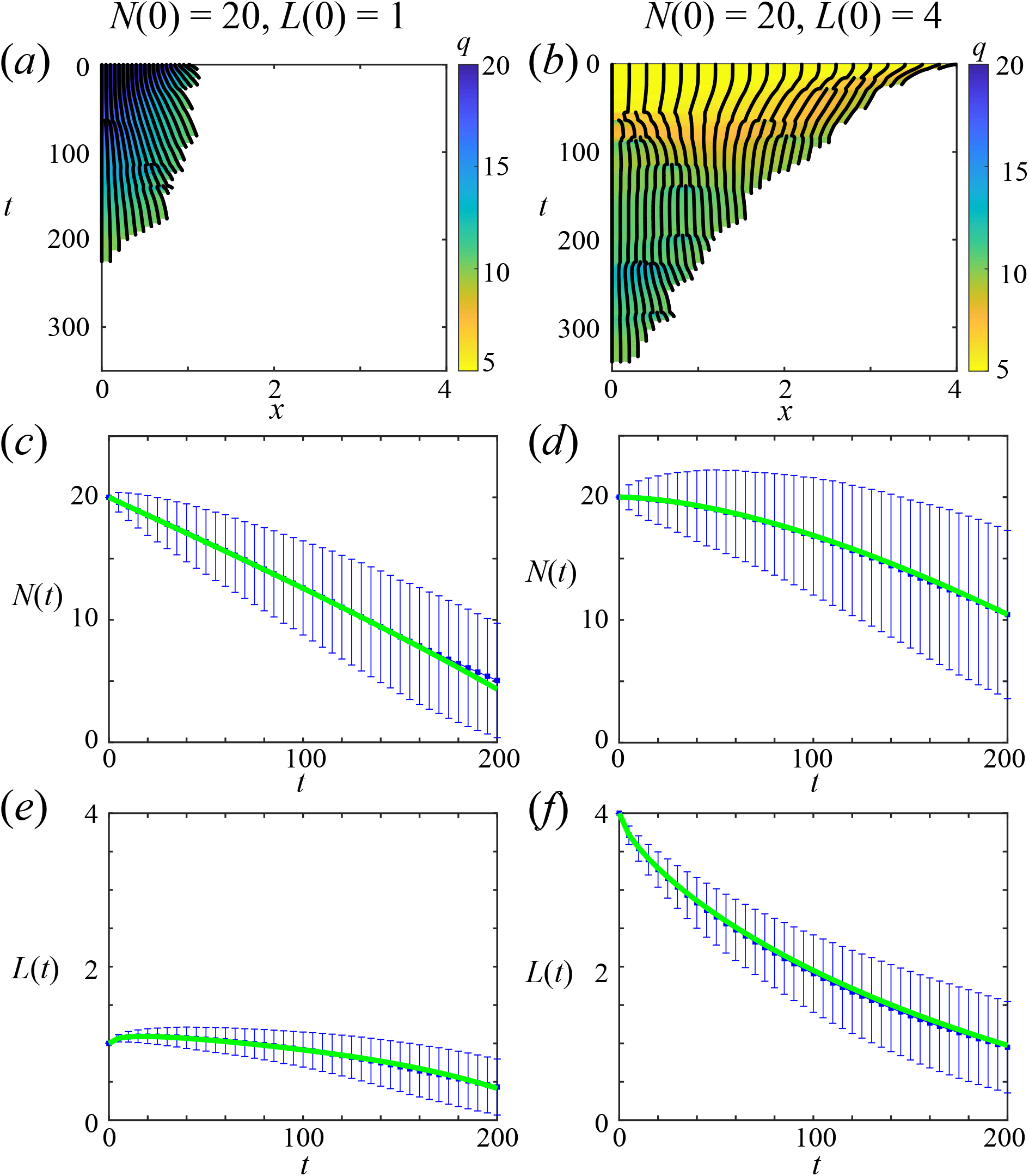
Chemically-independent cell detachment at a constant rate with a linear cell-length-dependent proliferation mechanism. Two initial cell populations with *N* (0) = 20, the first uniformly compressed with *L*(0) = 1 and the second uniformly stretched with *L*(0) = 4. (a)-(b) Kymographs with density, *q*(*x, t*), colouring. (c)-(f) The average of 2000 discrete realisations (blue) are compared with the continuum model (green). (c)-(d) Evolution of total cell number, *N* (*t*). (e)-(f) Evolution of tissue length, *L*(*t*). Mechanical parameters: *k* = 1, *a* = 0.1, *η* = 1. Critical tissue length is *ωa/β* = 4.

When we consider cell-length-dependent proliferation, the long term outcome of the model depends upon the mechanical properties, *k/η*, and rate of cell proliferation and detachment. In general, when *k/η* is large compared to *β*, the outcome of the model is similar to the simpler cell-length-independent proliferation case. As before, the solution of the continuum model is a good approximation of appropriately averaged data from the discrete model, except when *N* (*t*) is low (Figure 5). However, whereas for cell-length-independent proliferation stochastic effects are important when the initial number of cells *N* (0) is close to *ω/β*, for cell-length-dependent proliferation stochastic effects are important whenever the current tissue length, *L*(*t*), is close to the critical tissue length, *ωa/β*, as the epithelial tissue may go extinct in some realisations while the epithelial tissue may grow in other realisations (Appendix D.3). As we are considering chemically-independent EMT the total number of cells which detach grows linearly with time at rate *ω* when the tissue is not close to extinction (Figure 10).

### 3.3 Chemically-dependent EMT with small diffusivity

We now consider a general EMT-inducing chemical, with TGF-*β* being one such example of many candidate signalling molecules. As different EMT-inducing chemicals may have different diffusivities, we will consider a range of possible diffusivities from a very small diffusivity to assuming the chemical in the tissue is in diffusive equilibrium at all times. To begin, we assume very small diffusivity and cell-length-independent proliferation. Assuming that the tissue initially contains no chemical, the chemical is provided to the boundary cell only, and small diffusivity, then the chemical is mostly concentrated in the boundary cell. To compare the chemically-dependent cell detachment model with the chemically-independent cell detachment model (Section 3.1) we choose parameters so that the average rate of cell detachment is the same in both models, provided the boundary cell is close to its resting cell length. Chemically-dependent cell detachment is a two-step process: i) the boundary cell gains an invasive phenotype when its chemical concentration is above the chemical threshold, *C*, ii) the boundary cell detaches. So we introduce a parameter *ϕ* ∈ [0, 1] which defines the ratio of the average time in process i) as *ϕ/ω* and the average time in process ii) as (1 − *ϕ*)*/ω* (Figure 2). Note that *ϕ* = 0 corresponds to the chemically-independent model we explore in Sections 3.1-3.2. As before, the total number of cells which detach grows linearly with time at rate *ω* when the tissue is not close to extinction (Figure 10).

We find that agreement between results from the discrete model and corresponding continuum model is not as accurate as before for large values of *ϕ* (Figures 6(a),(d),(g) for *ϕ* = 0.9, Appendix D.5.1). In the discrete model we assume that a constant number of molecules of the chemical are supplied to the boundary cell (Equation (6)), to mimic a chemical diffusing in from the external environment, and assume that the concentration in every cell is well-mixed. However, for low diffusivities, here *D* = 10^−5^, the well-mixed assumption is not valid. So the rate of cell detachment, *ω*(*c*_*N*_ (*t*)), should be updated to account for intracellular chemical concentration gradients. In contrast to the discrete model, the continuum model does capture intracellular concentration gradients and the rate of cell detachment is determined by *c*(*L*(*T*), *t*), which is the concentration at the right edge of the boundary cell. With low diffusivity, the chemical concentration is localised to *L*(*t*) in the continuum model, rather than spread throughout the cell, so the continuum model reaches the concentration threshold faster than realisations of the discrete model. This explains the difference in Figures 6(d),(g). When *ϕ* is small or diffusivity is increased these differences are smaller (Appendix D.6,3.4).

**Figure 6.**
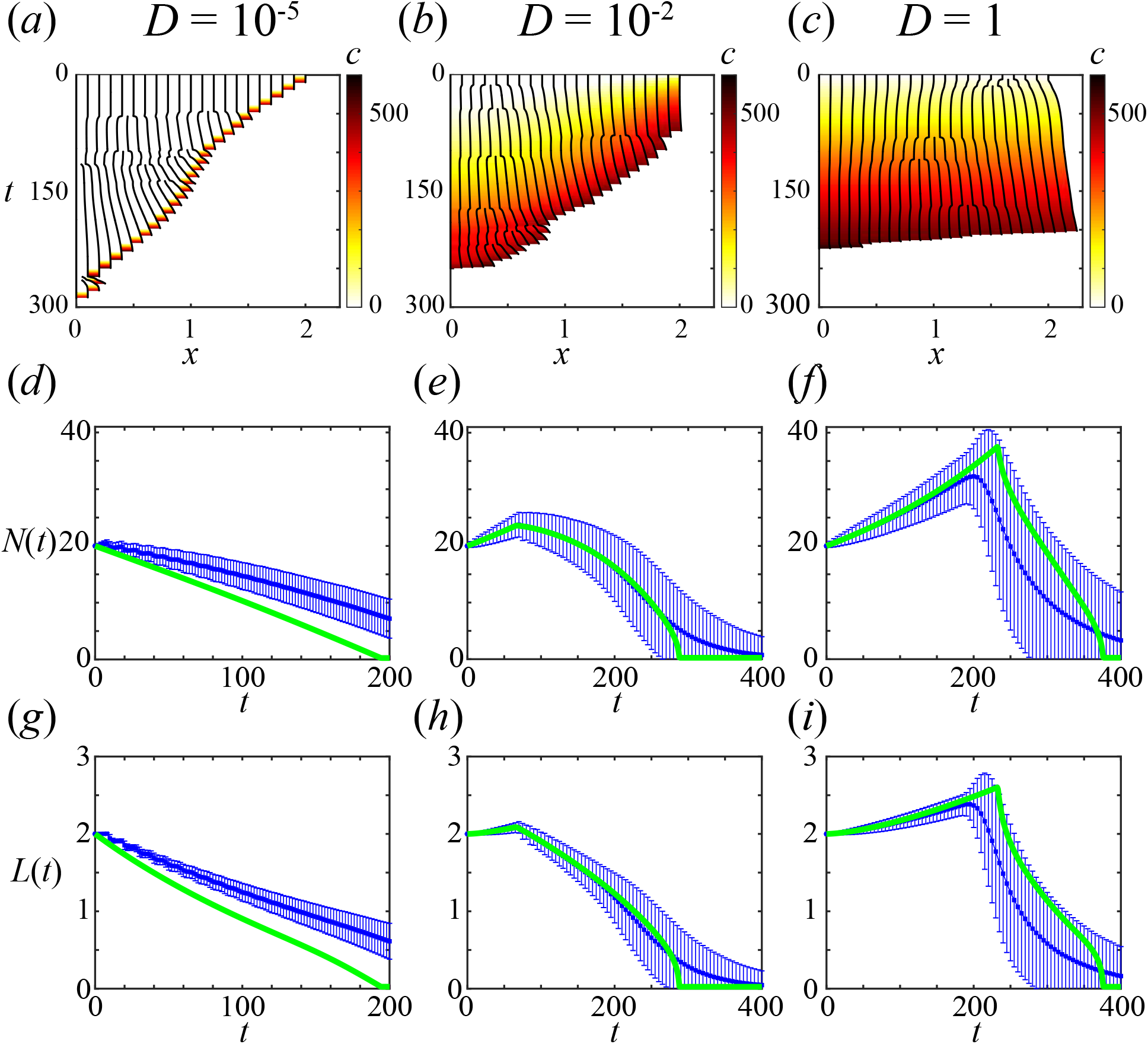
Increasing diffusion delays first EMT. Cell detachment driven by chemically-dependent EMT with varying diffusivities and a cell-length-independent proliferation mechanism. Cells are initially at their resting cell lengths with initial cell populations *N* (0) = 20. Kymographs with chemical concentration, *c*(*x, t*), colouring shown for (a) *D* = 10^−5^, (b) *D* = 10^−2^, (c) *D* = 1. (d)-(i) The average of 2000 discrete realisations (blue) are compared with the continuum model (green). (d)-(f) Evolution of total cell number, *N* (*t*). (g)-(i) Evolution of tissue length, *L*(*t*). Mechanical parameters: *k* = 1, *a* = 0.1, *η* = 1.

### 3.4 Chemically-dependent EMT with higher diffusivity

Higher chemical diffusivity results in the boundary cell having a lower chemical concentration, on average, than the same simulation with lower chemical diffusivity. This means that the time for the first cell to undergo EMT and detach increases (Figures 6(b),(c)). The delay of the first cell detachment can be sufficient to result in a transient rise in total cell number. However, as diffusivity is high the chemical concentration inside internal cells is close to the chemical concentration inside the boundary cell. Therefore, after the first cell detaches the new boundary cell may already be above or close to the concentration threshold, and hence quickly gains the invasive phenotype. This can lead to a rapid sequence of cell detachments, which was not seen with the models in the previous sections (Figures 6(b),(c), 10(e)-(g)).

Results in Figures 6(e),(f),(h),(i) show good agreement between the continuum model and the appropriately averaged quantities from many discrete realisations. The difference in Figures 6(e),(h) for *t* ≥ 250 is due to low *N* (*t*) near extinction (Section 3.1, Appendix D.4). The difference in Figures 6(f),(i) at *t* ≥ 200 is due stochastic effects in the discrete model, including the number of cells and tissue length. These are more prominent for *D* = 1 (Figures 6(f),(i)) in comparison to *D* = 10^−3^ (Figures 6(e),(h)) due to the increased time to reach the concentration threshold (Appendix D.5.2).

Special cases assuming the chemical in the tissue is at diffusive equilibrium at all times and instantaneous mechanical relaxation result in four possible behaviours (Appendix E): unbounded tissue growth without EMT and cell detachment; unbounded tissue growth with some EMT; eventual tissue homeostasis and constant EMT; and eventual tissue extinction due to EMT.

## 4 Conclusion

In this work we seek to explore how mechanical interactions influence the evolution of an epithelial tissue. Using our mathematical model, we ask when do tumours grow or shrink, and how mechanical interactions and an EMT-inducing chemical could influence the rate of cell detachment. Starting with a stochastic cell-based discrete model describing mechanical relaxation, cell-length-dependent proliferation, and cell detachment driven by chemically-dependent EMT, we derive the corresponding deterministic continuum description which takes the form of a novel nonlinear free boundary problem. In contrast to previous free boundary models we derive the boundary condition from cell-level biological processes and incorporate EMT. Both the discrete and continuum models useful information: discrete models show the important role of stochastic effects while continuum models help classify possible behaviours. Our results show good agreement between the continuum model and appropriately averaged quantities from many discrete realisations. However, as can be expected, there are occasions when the deterministic continuum model does not capture the fact that, due to stochastic proliferation and EMT in the discrete model, different identically prepared individual discrete realisations may result in different long-term behaviour.

Our models suggest that the coupling of mechanical interactions with EMT is important, can change the probability of long-term extinction significantly, and give rise to different rates of cell detachment [16, 17]. Using our model we postulate that to prevent cell detachment driven by EMT and delay the start of the metastatic cascade, one could chemically alter the speed of mechanical relaxation to encourage the tissue length to increase and hence cause the EMT-inducing chemical concentration to decrease. However, if the tissue length increases then proliferation may be more likely and the number of cancer cells in primary tumour would increase, which is also not desirable. Therefore, there is a delicate trade off between proliferation and EMT that should be considered when seeking to prevent cancer development. Furthermore, the model predicts that if EMT is delayed then the tissue may rapidly collapse due to many cells detaching in quick succession, which is undesirable as it may encourage metastasis. In contrast, for wound healing we may prefer cell-detachment driven by EMT and more proliferation to encourage the wound to heal faster. It will be useful to explore these ideas by extending this model to track individual cells or clusters of cells [19] that detach from the tissue in a two-regime model [49] or multi-organ model [50, 51], and incorporating mesenchymal-epithelial transitions (MET) [52, 53, 54]. Furthermore, the time taken for a cell to proliferate can be on the order of once every 12 hours to once every few days and EMT can occur over the course of hours, a few days [46], or many days (e.g. 9-12 days [47]), depending on the cell type and driving mechanisms. Therefore, the critical cell number *ω/β* and critical tissue length *ωa/β* µm we consider in this work are of a similar order of magnitude to that expected *in vitro* and may be interesting to test experimentally.

The mathematical framework that we develop here, and in related studies [30, 31, 32, 35, 36], is well-suited to incorporate additional biological mechanisms and explore different modelling assumptions. One modelling assumption we could change would be to allow the EMT-inducing chemical to diffuse across the boundary at the rear of the tissue, which may prevent a build up of chemical. Introducing intracellular diffusion in the discrete model would also resolve the issue around the well-mixed assumption not being valid for low diffusivities. We also assume, to illustrate a potential role for mechanochemical coupling, that a single chemical drives EMT. In reality, many biological processes at the level of proteins, mRNAs, and miRNAs occur and it may be interesting to incorporate regulatory networks which govern these processes into each cell in the discrete model [12, 13]. While linear diffusion of the EMT-inducing chemical between cells is arguably the simplest approach to include chemical transport and some experimental evidence exists for intercellular chemical transport [55], other mechanisms may be more biologically realistic, such as chemicals diffusing externally and being uptaken by cells [18] and cell adhesion regulated by interactions between E-cadherin and *β*-catenin [17]. It is an interesting question to ask whether other transport mechanisms are well approximated by the linear diffusion model we consider here.

The one-dimensional approach taken in this work has many advantages in its predictive power, interpretability, and relative computational simplicity in comparison to two- or three-dimensional models. Furthermore, cell-length-dependent proliferation may be thought of as an approximation for cell-volume-dependent proliferation which occurs for cells that move in three-dimensional environments. However, real cells can also spread without changing volume, so it may be beneficial to explore the role of the cell cycle in this one-dimensional framework [56]. A significant extension of this work would be to consider higher dimensions. The discrete model could be extended by considering a cell-centre or vertex model which introduces questions regarding cell shape and how neighbours can be identified, along with increased computational expense [39, 57, 58]. A corresponding continuum model in higher dimensions is less clear. The one-dimensional model enforces an ordering of neighbouring cells, which is important when deriving a continuum model [33, 59]. However, in higher dimensions cells can change their neighbours which poses significant challenges [33, 59]. We leave this extension for future consideration.

## Data Access

Key algorithms used to generate results are available on [Github](https://github.com/ryanmurphy42/Murphy2020b).

## Author Contributions

Authors R.J.M., P.R.B., R.E.B., and M.J.S. conceived and designed the study; R.J.M. performed numerical simulations and drafted the article; E.W.T. and H.J.H. provided advice on cancer and EMT; all authors provided comments and gave final approval for publication.

## Competing Interests

We have no competing interests.

## Funding

This work was funded by the Australian Research Council (DP170100474, DP200100177). R.E.B is a Royal Society Wolfson Research Merit Award holder, and also acknowledges the BBSRC for funding via grant no. BB/R000816/1. The Translational Research Institute receives funding from the Australian Government.

## A Continuum model: Evolution of free boundary equation derivation

The discrete model allows for cells to detach at the free boundary, *x* = *L*(*t*). In the continuum model this corresponds to loss of tissue material at a moving interface [43]. By considering conservation of mass for the total number of cells, *N* (*t*), the rate of change of *N* (*t*) due to proliferation and invasion is

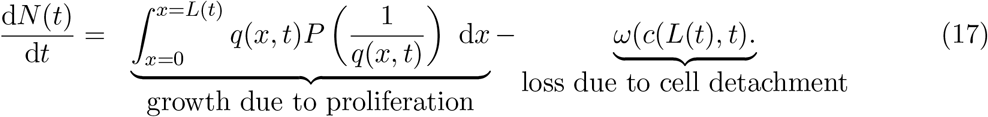

The rate of change of *N* (*t*) can also be written in terms of the cell density, *q*(*x, t*), as follows

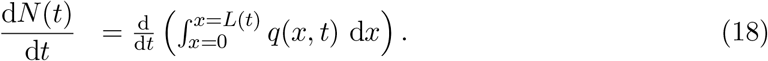

Differentiating the right hand side of Equation (18) with respect to *t* and applying Equation(8) for the cell density and Equation (9) for the boundary condition at *x* = 0 gives

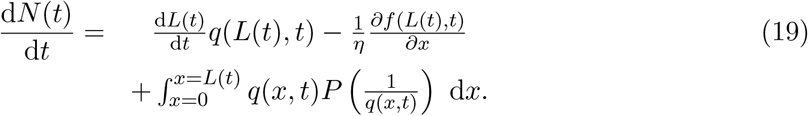

Equating (17) and (19) and rearranging we obtain evolution of the free boundary equation

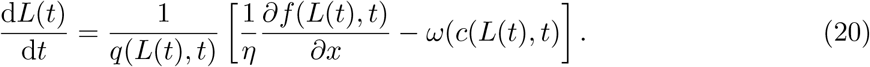

Substituting Equation (10) into Equation (20) we can obtain a different form for the evolution of the free boundary equation as

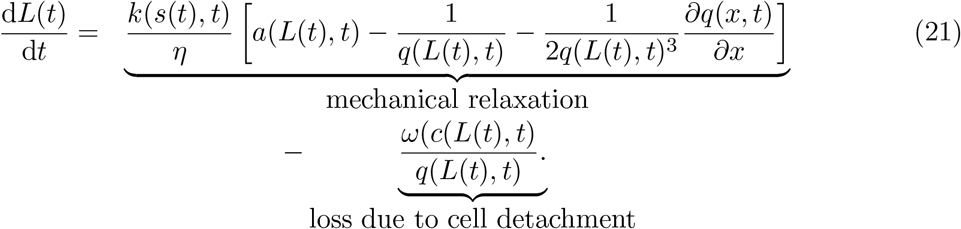

## B Numerical methods: Discrete model

The key technical challenge to overcome to numerically simulate the discrete model concerns how to implement linear chemical diffusion in an evolving population of cells with variable cells lengths. This is the primary focus of the first half of this section. Previous models have implemented diffusion on growing domains [24, 25, 26,27, 28, 29, 41, 44], however, what is unique to this work is that we are interested in the chemical concentration inside individual cells when the positions of cell boundaries are known and evolve in time. Furthermore, previous studies tend to consider uniform growth throughout the tissue whereas here, due to mechanical interactions and proliferation, we have non-uniform growth throughout the tissue. As we will show these complications requires a new numerical method.

To model the diffusion of molecules of a chemical we have a choice between microscopic or mesoscopic individual-based models or macroscopic population-based models. Microscopic individual-based models are often posed as a population of particles undergoing Brownian motion. Mesoscopic individual-based models are often posed as a population of particles undergoing a random walk on a lattice or as a position-jump process on a lattice. However, individual-based models tend to be more computationally expensive than population-level models and can be mathematically intractable. Macroscopic approaches can be simpler to write down and are often easier and faster to simulate for a large number of particles.

Macroscopically, linear diffusion of particles on a fixed domain, 0 < *x* < *L*, can be modelled with the following classical partial differential equation

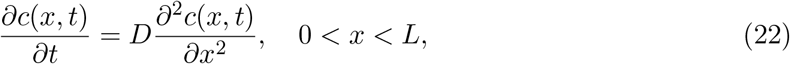

where *c*(*x, t*) is the particle density, or equivalently the chemical concentration, and *D* is the macroscopic diffusion coefficient. Whereas on a domain whose length, *L*(*t*), evolves in time, conservation of mass arguments and applying Reynolds transport theorem gives the following partial differential equation for the evolution of *c*(*x, t*), [44]

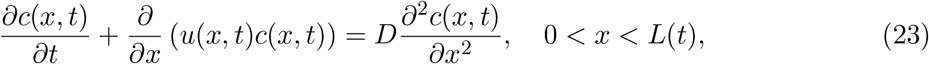

where *u*(*x, t*) is a velocity field prescribed by domain growth [24, 25, 41, 44]. In this work, Equation (23) is the same as Equation (13) in the main manuscript with *u*(*x, t*) = 1*/*(*ηq*(*x, t*)) *∂f* (*x, t*)*/∂x* determined by mechanical interactions between the cells.

Previous studies have demonstrated that a stochastic individual-based model incorporating domain growth, taking the form of a position-jump model on a uniform lattice, is equivalent to the continuum model in Equation (23) [44]. However, domain growth was implemented by instantaneous doubling and dividing of underlying lattice sites which always results in a uniform lattice, corresponding to all cells have the same length at all times. This is not the case for this work. We obtain a non-uniform lattice, as cell lengths vary due to the effects of mechanical interactions between cells and proliferation. Therefore, to discuss the method we will focus on a non-uniform lattice and initially assume that positions of the cell boundaries are fixed and known.

To model diffusion on a non-uniform lattice Yates et al. [41] make clear that one must be careful, and suggest two methods which we refer to as method A and method B. To explain the methods we first state the two key sets of points: i) positions of cell boundaries, *x*_*i*_ for *i* = 1, 2, …, *N* + 1 (represented as circles in schematics in Figures 1,7-9); ii) resident points, *y*_*i*_ for *i* = 1, 2, …, *N*, satisfying *x*_*i*_ < *y*_*i*_ < *x*_*i*+1_, which defines the location in cell *i* where the particles of the chemical are considered to be positioned (represented as crosses in schematics in Figures 7-9). In method A (Figure 7(a)), Yates et al. [41] assume the resident points, *y*_*i*_, are chosen first and then the cell boundaries, *x*_*i*_, are defined in a Voronoi neighbourhood sense: a point is in cell *i* if it is closer to the resident point associated with cell *i*, given by *y*_*i*_, rather than any other resident point *y*_*j*_. They show this method can be used to accurately model linear diffusion due to the Voronoi partition (see Supplementary Material Section 1 of Yates et al. [41]). However, in this work the positions of the cell boundaries are already known from the mechanical interactions (Equations (1)-(3)) so we cannot use method A and instead consider method B. In method B (Figure 7(b)), Yates et al. [41] first prescribe the position of the cell boundaries, *x*_*i*_, which is what we require, and then they choose the resident points to be the position of the centre of cell *i*, so *y*_*i*_ = (*x*_*i*_ + *x*_*i*+1_)*/*2. They show this method does not accurately model linear diffusion as there is not a Voronoi partition. When all cells are the same size, resulting in a uniform lattice, methods A and B are equivalent.

**Figure 7.**
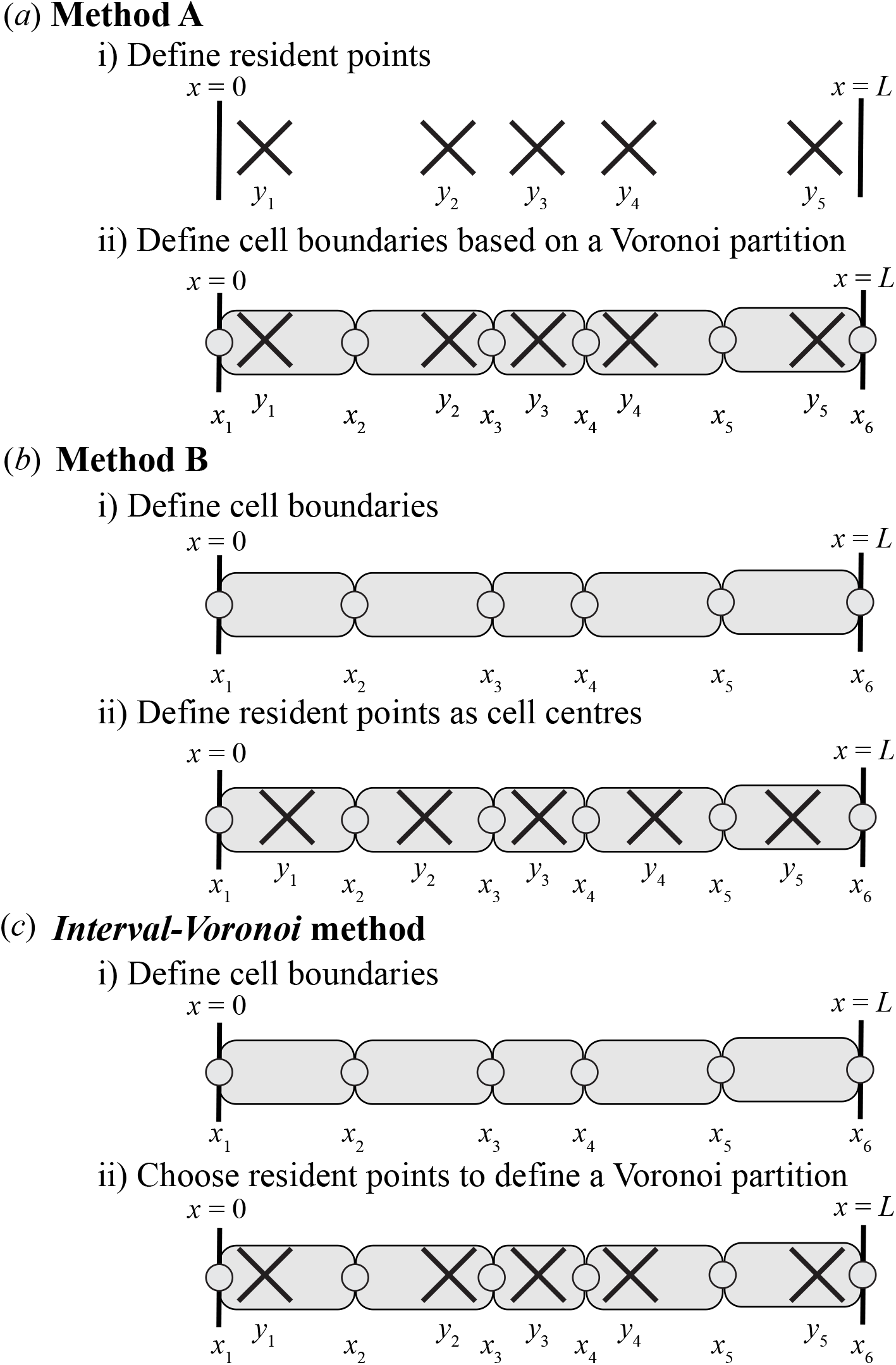
Different methods to model linear diffusion for variable cells sizes. (a) Method A: define resident points, *y*_*i*_, first and then define cell boundaries, *x*_*i*_, to form a Voronoi partition. (b) Method B: define cell boundaries first and then define resident points at the cell centres. (c) New Interval-Voronoi method: define cell boundaries first and then choose resident points to define a Voronoi partition. Circles represent cell boundaries, *x*_*i*_, and crosses represent resident points, *y*_*i*_. Shown for *N* = 5 cells.

Before proceeding with our new method, which combines and extends methods A and B, we briefly discuss the underlying microscopic diffusion model and how it relates to the mesoscale position-jump process model of diffusion [41]. In the microscopic model of diffusion, the position of an individual particle which undergoes Brownian motion is governed by a stochastic differential equation,

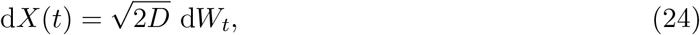

where d*W*_*t*_ is a standard Wiener process and *D* is the macroscopic diffusion coefficient. For the mesoscale position-jump process model of diffusion we seek the rates at which a particle at resident point, *y*_*i*_, moves to the neighbouring left or right resident points at *y*_*i*−1_ or *y*_*i*+1_, respectively. These transition rates, 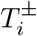, can be found by initialising a particle at *y*_*i*_ solving a first passage time problem on the domain *y*_*i*−1_ < *x* < *y*_*i*+1_ [41, 60]. Then the transition rates are [41, 61]

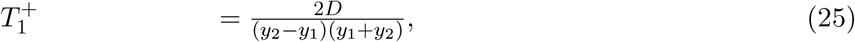

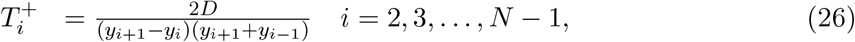

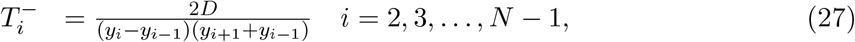

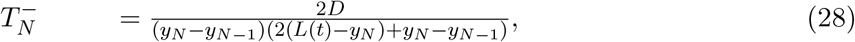

where *D* is the macroscopic diffusion coefficient. When all cells are the same length, *l*, these transition rates simplify to *D/l*^2^.

To accurately model linear diffusion in our work we combine and extend methods A and B. Specifically, we first define the positions of the cell boundaries, *x*_*i*_, and then we choose the resident points, *y*_*i*_, so that the position of the bisection of neighbouring resident points is the position of a cell boundary, i.e. choose *y*_*i*_ such that (*y*_*i*_ + *y*_*i*+1_)*/*2 = *x*_*i*_ for *i* = 2, 3, …, *N* (Figure 7(c)). This results in a Voronoi partition on the set of *y*_*i*_, where the edges of the Voronoi partition coincide with the cell boundaries. We now explain how to choose the *y*_*i*_ in such a manner by following Figure 8. We assume an initial cell configuration (Figure 8(a)) and will work from the leftmost cell to the rightmost cell. Initially, the resident point of the leftmost cell could be placed anywhere in this cell, which we call the possible region of the first cell and show in green (Figure 8(b)). Next we reflect the possible region of the first cell about the right boundary of the first cell, and colour this yellow (Figure 8(c)). Then intersecting the reflected possible region of the first cell with the interval occupied by the second cell gives the possible region for the second cell, which we indicate in green (Figure 8(d)). We repeat until reaching the rightmost cell (Figure 8(e)-(h)). Given possible regions for all of the cells we choose the resident point of the rightmost cell to be the midpoint of the possible region of the rightmost cell (Figure 8(i)). Then reflecting the resident point about the left boundary of the rightmost cell we obtain the location of the resident point of the penultimate cell. We repeat until we have obtained the resident points for all cells (Figure 8(j)). This method gives us a set of resident points which can be used to accurately model linear diffusion. However, this method only works when a Voronoi partition can be defined, which occurs when cells are similar lengths.

**Figure 8.**
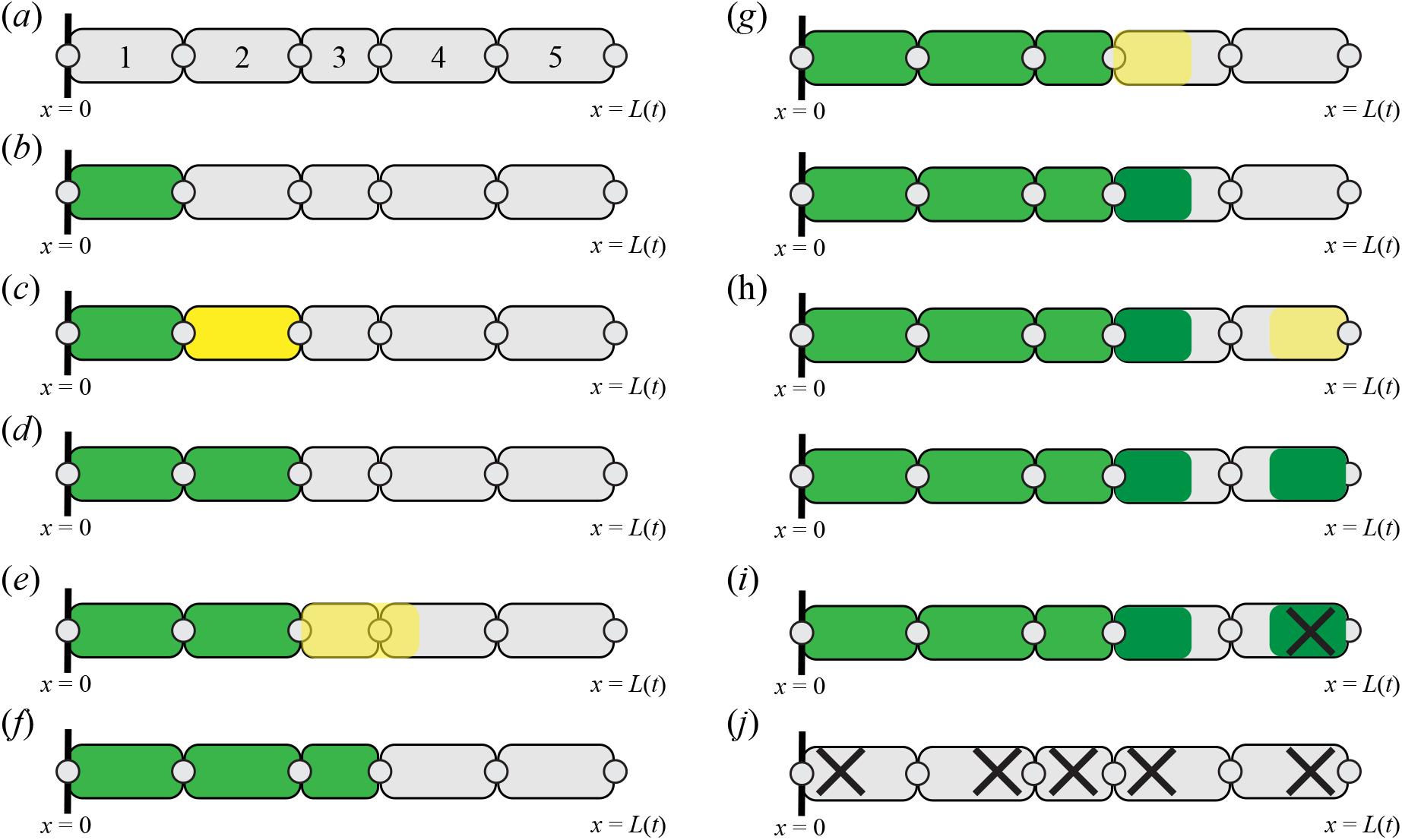
Schematic for *Interval-Voronoi* method for cells of similar size. Circles represent cell boundaries, *x*_*i*_, crosses represent resident points, *y*_*i*_, and green represent possible regions for a cell where a resident point could be placed. Details discussed in the text.

Depending on the configuration of the positions of the cell boundaries it is not always possible to define the resident points as above. For example, following Figure 9, the possible regions for the first three cells can be defined as before (Figure 9(a)-(f)). However, when we reflect the possible region of the third cell and intersect this with the interval occupied by the fourth cell we obtain an empty set. Therefore, there is no location in the third cell where we will be able to place a resident point to define a Voronoi partition. To resolve this problem, we now divide the third cell into two compartments (Figure 9(h)). To do so, we first assume that the current possible region for the third cell is now the possible region for the left compartment of the third cell. Then, we choose the position of the compartment boundary which divides the cell (red-dashed line in Figure 9(h)) so that the position of the right boundary of the right compartment of the third cell is equal to the position of the right boundary of the third cell, the possible region of the left compartment of the third cell does not overlap with the possible region of the right compartment of the third cell, and the possible region of the right compartment of the third cell is maximised. In Figure 9(h) this corresponds to dividing the cell into two compartments of equal length. We can then proceed as before (Figure 9(i)-(j)). We can then define resident points for each compartment and subsequently for each cell (Figure 9(k)). When numerically simulating the discrete model, as described further below, we update the chemical concentration using a well-mixed assumption at each time step so this method does not result in intracellular chemical concentration gradients. This a valid approximation when *D* is sufficiently large.

**Figure 9.**
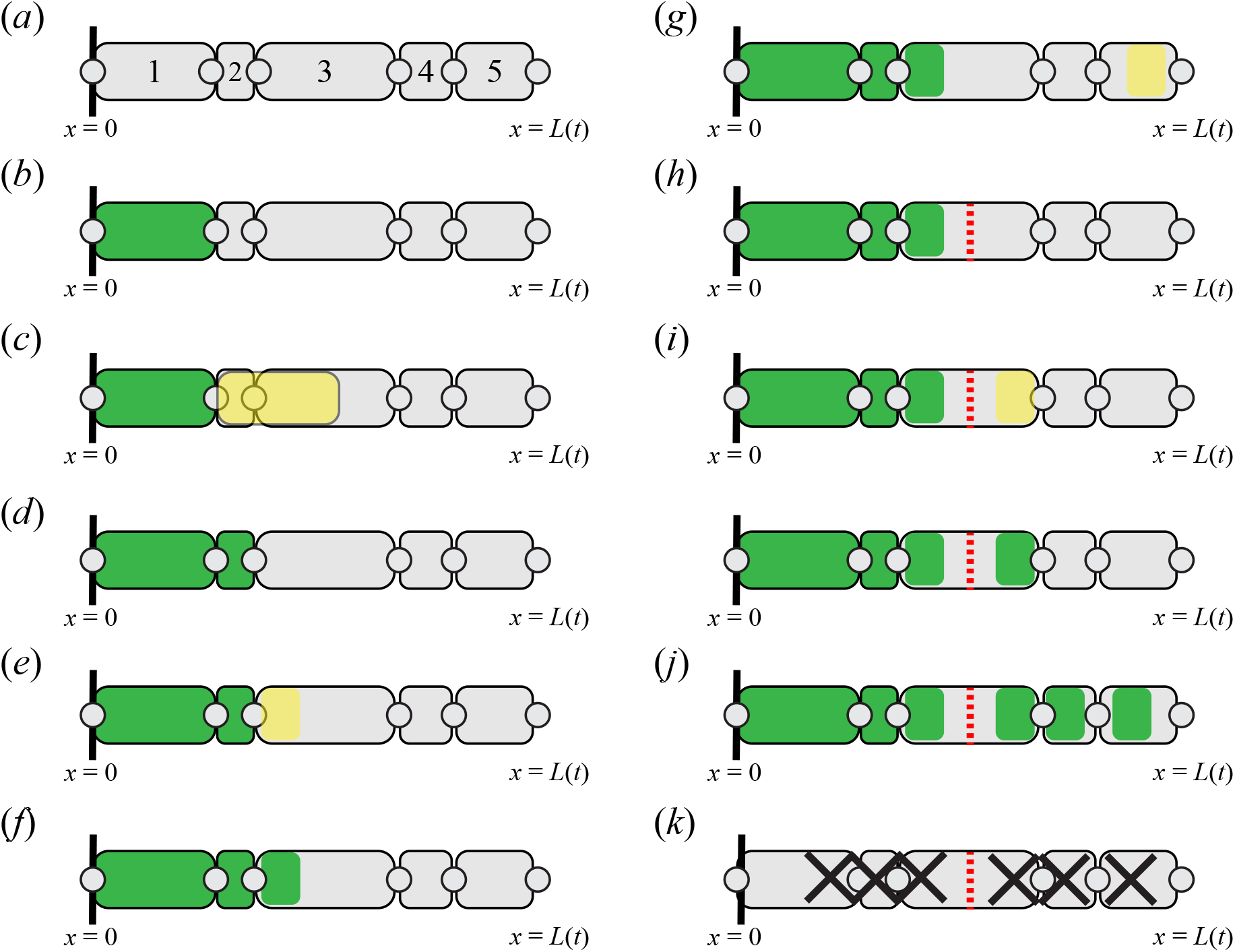
Schematic for *Interval-Voronoi* with compartments per cell. This method is required when a Voronoi partition cannot be defined on the initial cell configuration, which occurs when cells lengths are not similar. Circles represent cell boundaries, *x*_*i*_, crosses represent resident points, *y*_*i*_, green represent possible regions for a cell where a resident point could be placed, red-dashed represents the a compartment boundary within a cell. Details discussed in the text.

Being able to define a Voronoi partition is necessary [41] but results from simulations show that it is not always sufficient to accurately model linear diffusion. The distances between neighbouring resident points should also be the same order of magnitude throughout the population. If we suppose the resident points are *y*_*i*_ for *i* = 1, 2, …, *N* then if

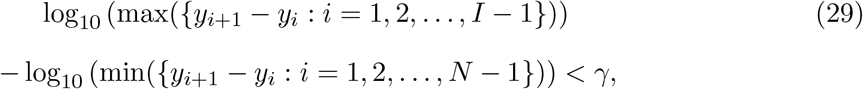

the distances between neighbouring resident points are approximately the same order and we can proceed. The condition in Equation (29) and the value *γ* = 0.8 were found to be good choices for results in this work. If the condition in Equation (29) is not satisfied we introduce a maximum compartment length equal to half of the minimum length of the current cell sizes. Then for any cells whose length is greater than the maximum compartment length, we divide those cells into compartments so that the maximum length of any compartment is less than the maximum compartment length. We then determine the resident points as above.

The method described above, which we name the *Interval-Voronoi* method, can now be used to accurately model linear diffusion on a fixed domain with variable cell lengths. In this work, all of the cell boundaries, and consequently the length of the domain, *L*(*t*), are evolving in time. However, when we numerically simulate a single realisation of the discrete model we discretise time with a constant time step Δ*t*. Then for each time interval [*t, t* + Δ*t*) we assume the cell boundaries are fixed and the domain is fixed and apply the *Interval-Voronoi* method. Further details are now shown.

Let us consider a single realisation of the discrete model. The epithelial tissue is initialised with *N* cells each with cell stiffness *k* and resting cell length *a*. The position of each cell boundary *x*_*i*_(*t*) for *i* = 1, 2, …, *N* is defined. Every cell in the tissue is prescribed with the same proliferation mechanism and proliferation parameter, *β*. The chemical concentration is initially set to zero in each cell. To simulate the model, we discretise time with a constant time step Δ*t*. Then for each time interval [*t, t* +Δ*t*) we: i) update cell positions according to mechanical interactions; ii) update the chemical concentrations in each cell; iii) implement proliferation or cell detachment if it occurs.

We update the positions of each of the cell boundaries, *x*_*i*_(*t*), by integrating Equations (1)-(3) using a forward Euler approximation,

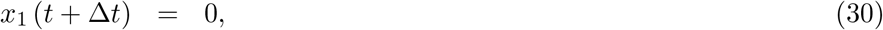

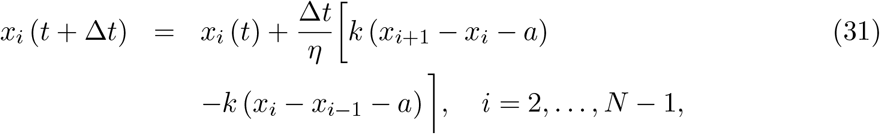

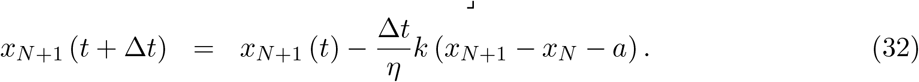

Next we seek to update the chemical concentration by applying the *Interval-Voronoi* method. Using the updated positions of cell boundaries, *x*_*i*_(*t* +Δ*t*) for *i* = 1, 2, …, *N* + 1, we determine the resident points *y*_*i*_ for 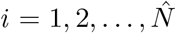. Then we can calculate the boundaries of compartments 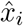 for 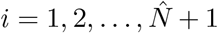, in a Voronoi neighbourhood sense, where 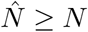 is the number of resident points. If the *Interval-Voronoi* method requires at least one cell to be divided into compartments then 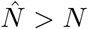 in order for a Voronoi partition to be defined. Then given the resident points we determine the transition rates, 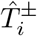, in terms of compartment lengths, 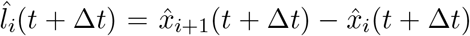 We now have the information required to update the chemical concentration for the cells. First we update the chemical concentration for all compartments, which we denote as *ĉ*_*i*_(*t*) for 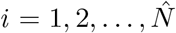 by integrating Equations (4)-(6) using forward Euler approximations,

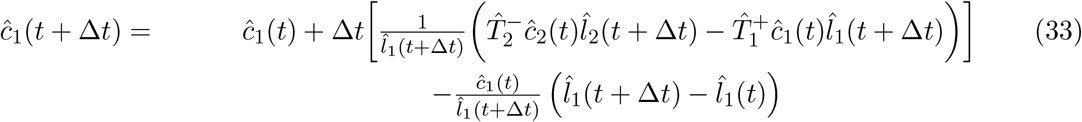

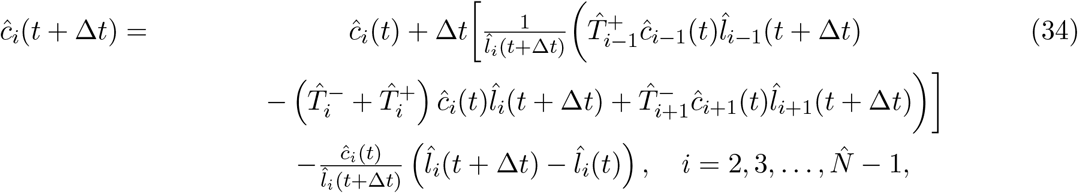

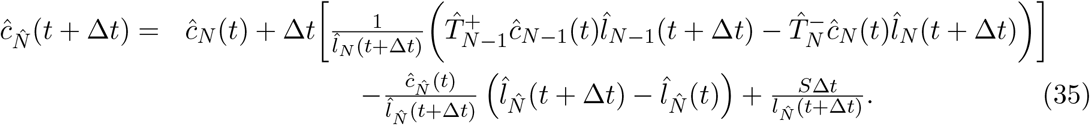

If the *Interval-Voronoi* method does not introduce any compartments per cell, i.e. if 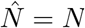, then 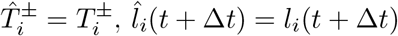, and *ĉ*_*i*_(*t*) = *c*_*i*_(*t*). Hence, if 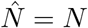, using Equations (34)-(35), we directly determine the chemical concentrations *c*_*i*_(*t* + Δ*t*) for *i* = 1, 2, …, *N* and we can proceed to incorporating if a cell proliferation or detachment event occurs in the time step. However, if 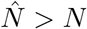 before proceeding we apply the well-mixed assumption to each cell. Specifically, if the *Interval-Voronoi* method introduces *j* compartments into cell *i* with concentrations *ĉ*_*k*_(*t* + Δ*t*), *ĉ*_*k*+1_(*t* + Δ*t*), …, *ĉ*_*k*+*j−*1_(*t* + Δ*t*) then the chemical concentration of cell *i* is set to be 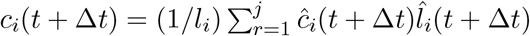.

Next, given the cell positions and chemical concentrations at time *t* + Δ*t*, we determine whether a cell proliferation event or a cell detachment event occurs during the time interval [*t, t*+Δ*t*). Analogous to Murphy et al. [36] we proceed using rejection sampling [62] where we generate three independent random numbers from a uniform distribution, *r*_1_ *∼ U* [0, 1], *r*_2_ *∼ U* [0, 1], *r*_3_ *∼ U* [0, 1]. Then a cell event, which could be either a proliferation event or a cell detachment event, occurs when

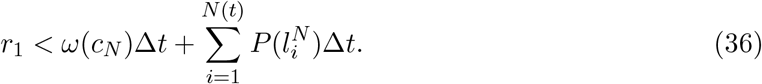

Given that a cell event occurs, a proliferation event occurs if

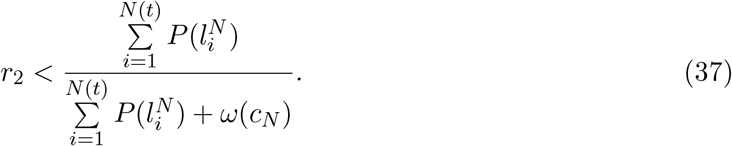

Otherwise we have a cell detachment event and we remove the boundary cell. If a proliferation event occurs, to determine which cell is proliferating we find the index *j* which satisfies

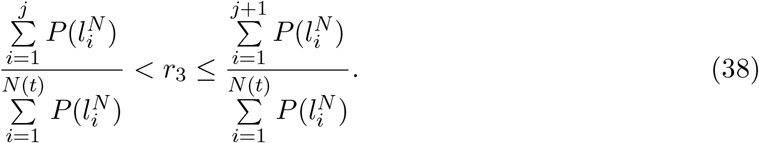

We then divide the parent cell into two equally sized daughter cells with the same chemical concentration and mechanical properties as the parent cell.

We repeat the above steps for each time step until the final time. This method requires that at most one cell event can occur within each time step so Δ*t* should be chosen sufficiently small.

### C Numerical methods: Continuum model

The continuum model is solved numerically using a boundary fixing transformation [45], finite difference approximations, and the Newton-Raphson method [63] with adaptive time stepping. Full details now follow.

For completeness, we rewrite the governing equations for the cell density, *q*(*x, t*), from Equation (8) and chemical concentration, *c*(*x, t*), from Equation (12), where we have used the definitions of *f* (*x, t*), and *u*(*x, t*) from Equation (13),

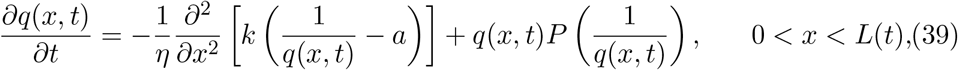

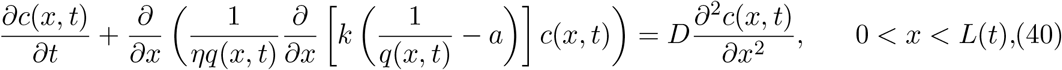

These governing equations are solved with the following boundary conditions from Equations (9), (10), (14), (15),

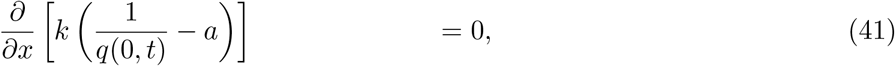

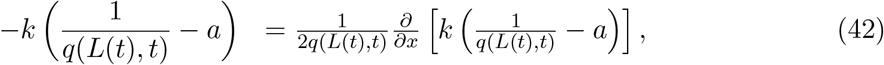

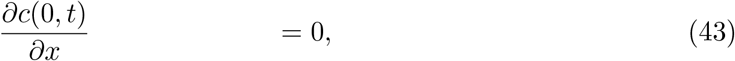

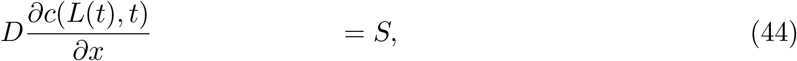

and the boundary position, *L*(*t*), evolves according to Equation (11) and we can apply Equation (42) to give

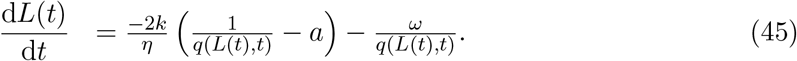

Equations (39)-(45) form a moving boundary problem for coupled non-linear partial differential equations. To proceed we apply a standard boundary fixing transformation [45] by setting *ξ* = *x/L*(*t*) to transform the evolving domain 0 *≤ x ≤ L*(*t*) to a fixed domain 0 *≤ ξ ≤* 1. Equations (39)-(45) then become

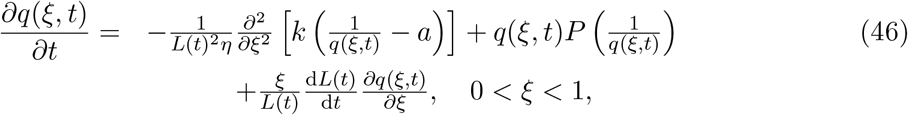

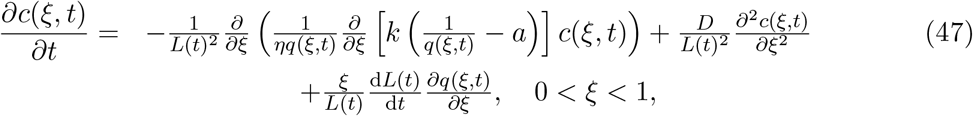

with the following boundary conditions

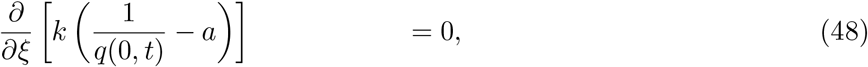

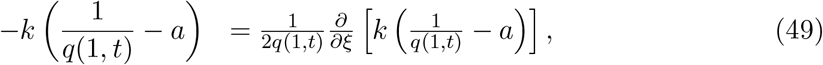

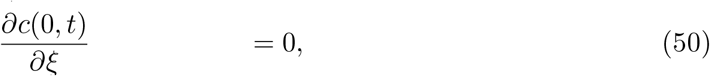

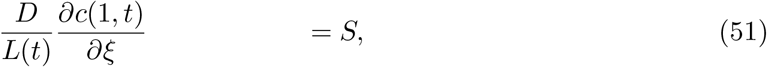

and the boundary position, *L*(*t*), evolves according to

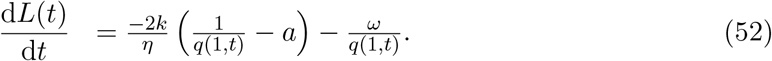

Next, we discretise the domain 0 *≤ ξ ≤* 1 with a uniform mesh with spatial step Δ*ξ* and use the subscript *j* = 1, 2, …, *J* to represent the index of the spatial nodes. We discretise time with a uniform mesh with time step Δ*t* and use the superscript *n* = 1, 2, …, *T* to represent temporal step. Second-order spatial derivatives are approximated by standard central differences. First-order spatial derivatives are approximated by standard upwind differences. A standard implicit finite difference approximation is used to approximate temporal derivatives.

Equation (52) governing the evolution of *L*(*t*) becomes

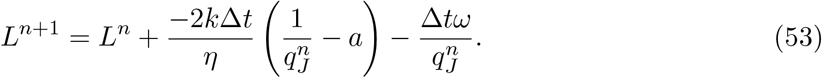

Finite difference approximations of the cell density equations give, for internal spatial nodes *j* = 2, …, *J −* 1,

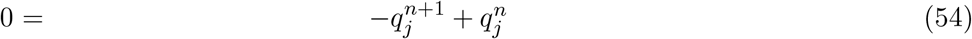

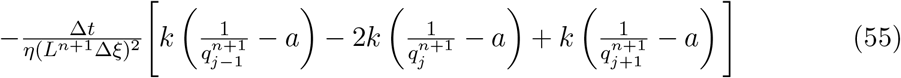

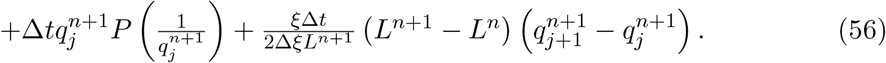

At *ξ* = 0, corresponding to spatial node 1, the cell density is updated with

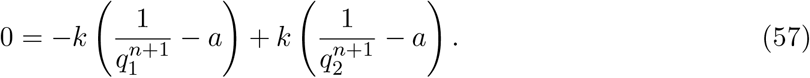

At *ξ* = 1, corresponding to spatial node *J*, the cell density is updated with

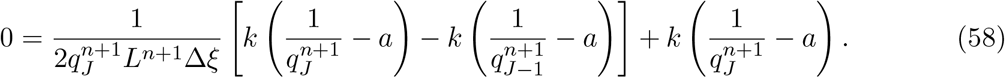

To proceed we calculate the velocity at each node, given by

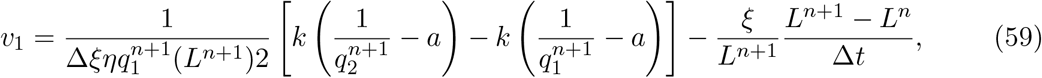

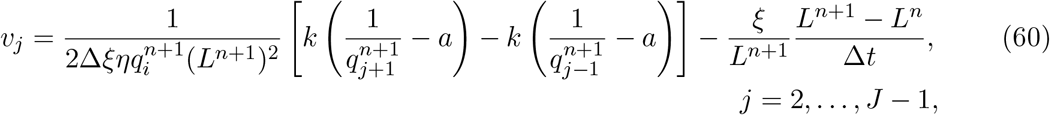

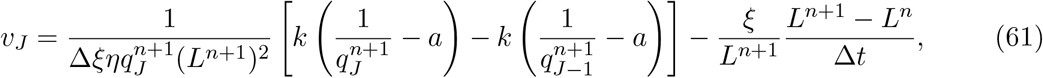

where forward and backward differences applied at the left and right boundaries, respectively. Considering the chemical concentration for internal nodes *j* = 2, 3, …, *J −*1 and assuming *v*_*j*_ *≥* 0 so that first order spatial derivatives are upwinded, finite difference approximations give

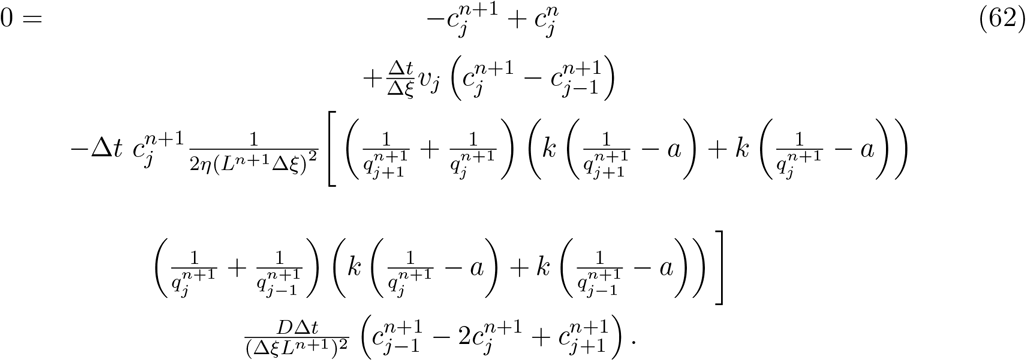

Similarly, if *v*_*j*_ *<* 0 upwinding is applied to the first order spatial derivative. At *ξ* = 0 we apply a forward difference approximation to Equation (50) governing the chemical concentration which gives

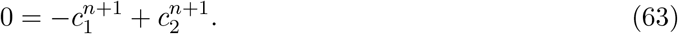

At *ξ* = 1 we apply a backwards difference approximation to Equation (51) governing the chemical concentration which gives

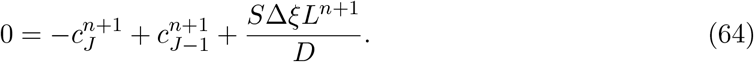

Equations (53)-(64) form a system of nonlinear algebraic equations for the cell density, chemical concentration, and evolution of *L*(*t*). We solve these equations using the Newton-Raphson method [36, 63]. In each Newton-Raphson iteration we first update *L*(*t*), according to Equation (53), then the cell density, governed by Equations (54)-(58), then the chemical concentration, governed by Equations (62)-(64). Newton-Raphson iterations are performed at each time point until the infinity norm of the the difference between successive estimates of 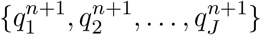 and 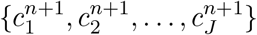 is below a specified tolerance,*ϵ*. To ensure that the Newton-Raphson iteration converges we apply adaptive time stepping. To implement adaptive time stepping we introduce a maximum number of iterations. When the maximum number of iterations is reached without the tolerance being met we divide the timestep by ten and repeat. Once the tolerance is met we reset the time step for the next temporal node. In our results adaptive time stepping is important when the chemical concentration first reaches the chemical threshold at *L*(*t*), and it reduces the computational time required to obtain the numerical solution. We use the Thomas algorithm to solve the linear systems which arise from the Newton-Raphson method. To ensure all numerical results are grid-independent we set Δ*ξ* = 10^*−*5^, initially set Δ*t* = 10^*−*3^, set the maximum number of iterations for each time step to ten, and set ε = 10^*−*8^.

Key algorithms used to generate results are available on [Github](https://github.com/ryanmurphy42/Murphy2020b).

#### C.1 Initial conditions

In this work, we refer to three types of initial conditions: compressed, mechanical equilibrium, and stretched. Here, we state these for the discrete and continuum model.

In the discrete model we choose every cell to initially have the same length, *l*_*i*_(0) = *L*(0)*/N* (0). If *l*_*i*_ *< a* then each cell is compressed and the tissue is compressed. If *l*_*i*_ = *a* then each cell is at its resting cell length and the tissue is at mechanical equilibrium. If *l*_*i*_ *> a* then each cell is stretched and the tissue is stretched. The corresponding initial conditions in the continuum model are obtained using *q*(*x*, 0) = *N* (0)*/L*(0) for 0 *< x < L*(0).

### D. Additional results

#### D.1 Counting the total number of cells that detach

We first define *M* (*t*) as the total number of cells which have detached by time *t*. For chemically-independent EMT, cell detachment occurs at a constant rate *ω*. Therefore, *M* (*t*) = *ωt* while *N* (*t*) *>* 0, i.e. the total number of cells which have detached increases linearly with time while the tissue still contains cells. If *N* (*t*) reaches zero then *M* (*t*) plateaus. In Figures 10(a)-(d), we show the good agreement between the results of the average of many discrete realisations and *M* (*t*) = *ωt*. This holds for later times in Figure 10(b) as the tissue does not go extinct. In contrast for Figures 10(a),(c),(d) *M* (*t*) eventually plateaus at later times due to extinction.

**Figure 10.**
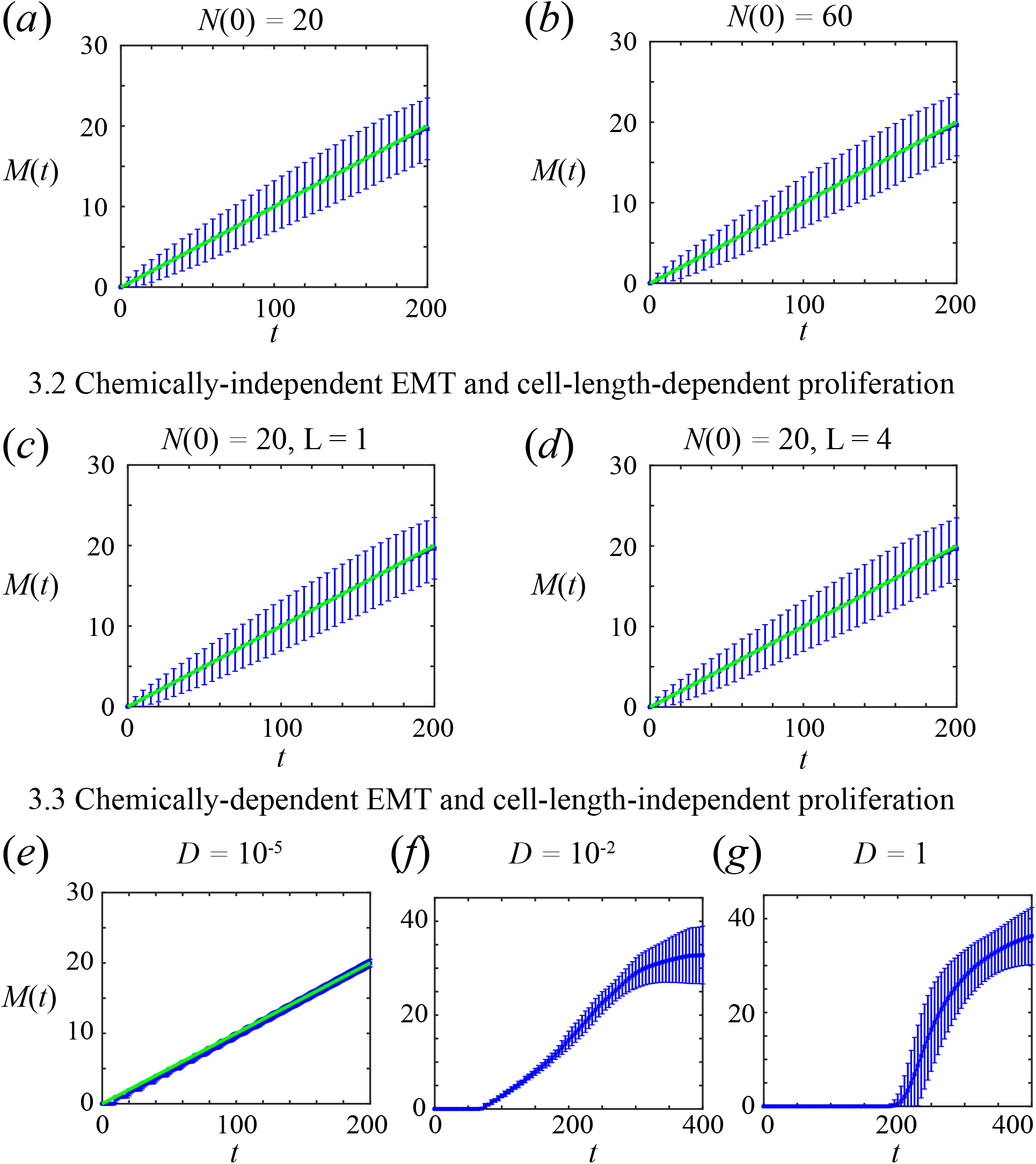
Evolution of total number of cells that detach, *M* (*t*), for examples presented in the main manuscript. (a)-(b) Chemically-independent EMT and cell-length independent proliferation (from Section 3.1). (c)-(d) Chemically-independent EMT and linear cell-length-dependent EMT (from Section 3.2). (e)-(g) Chemically-dependent EMT and cell-length-independent proliferation for (e) *D* = 10^*−*5^, (f) *D* = 10^*−*2^, (g) *D* = 1 (from Section 3.3-3.4). The average of 2000 discrete realisations (blue) are compared with *M* (*t*) = *ωt* (green).

For chemically-dependent EMT and *D* = 10^*−*5^ in Figure 10(e), *M* (*t*) is very similar to the chemically-independent EMT case due to how parameters are chosen, as discussed in Section 3.3. Also note that as *ϕ* = 0.9 in Figure 10(e) the cells require some time to reach the chemical threshold before detaching rapidly. Hence, we observe reduced noise for chemically-dependent EMT with *D* = 10^*−*5^ in comparison to results for chemically-independent EMT in Figures 10(a)-(d). As *ϕ →* 0 the noise in *M* (*t*) will increase, whereas for *ϕ* = 1 there will be no noise in *M* (*t*). For chemically-dependent EMT with higher diffusivities of *D* = 10^*−*2^ and *D* = 1, Figures 10(f),(g), respectively, cell detachment is initially delayed, then cells detach rapidly until extinction occurs.

### D.2 Cell-length-independent proliferation

In the main manuscript we present results for the evolution of *N* (*t*) and *L*(*t*) with *k* = 1 starting with cells initially at mechanical equilibrium. In Figures 12 and 13 we present results for *k* = 10 and tissues that are initially compressed or stretched. Good agreement is observed between the continuum model and the average of many discrete realisations.

**Figure 11.**
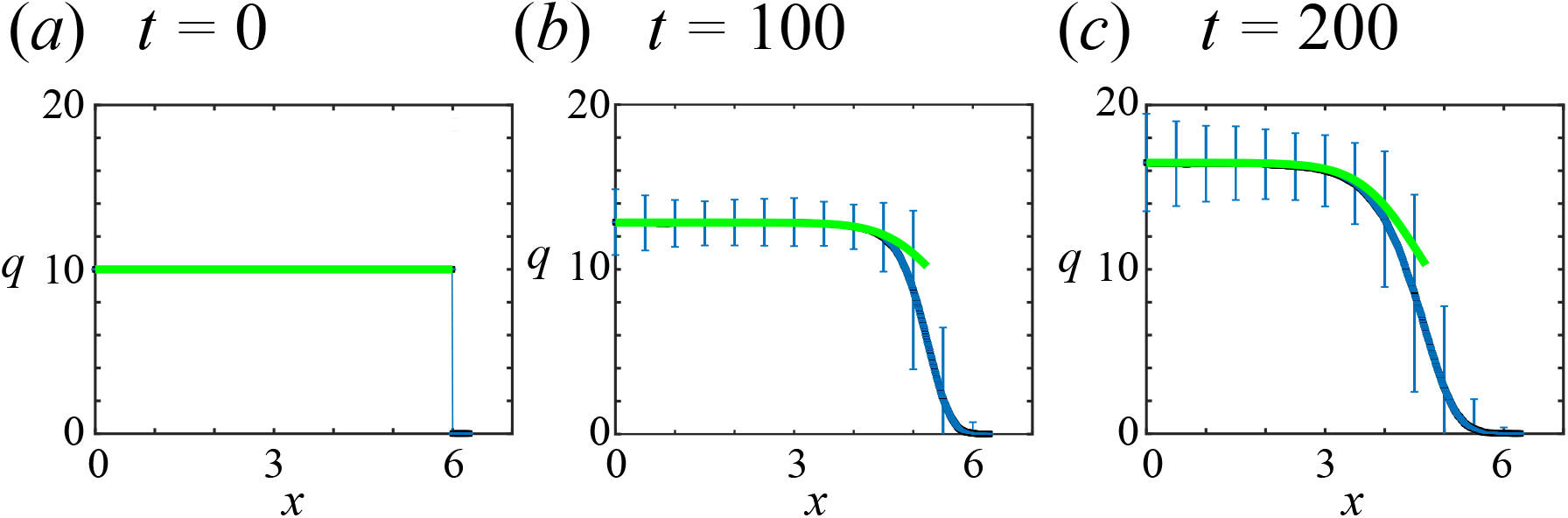
Density snapshots corresponding to *N* (0) = 60 in Figure 3. Cells are initially at their resting cell lengths. The average of 2000 discrete realisations (blue) are compared with the continuum model (green) at times (a) *t* = 0, (b) *t* = 100, (c) *t* = 200. Mechanical parameters: *k* = 1, *a* = 0.1, *η* = 1.

**Figure 12.**
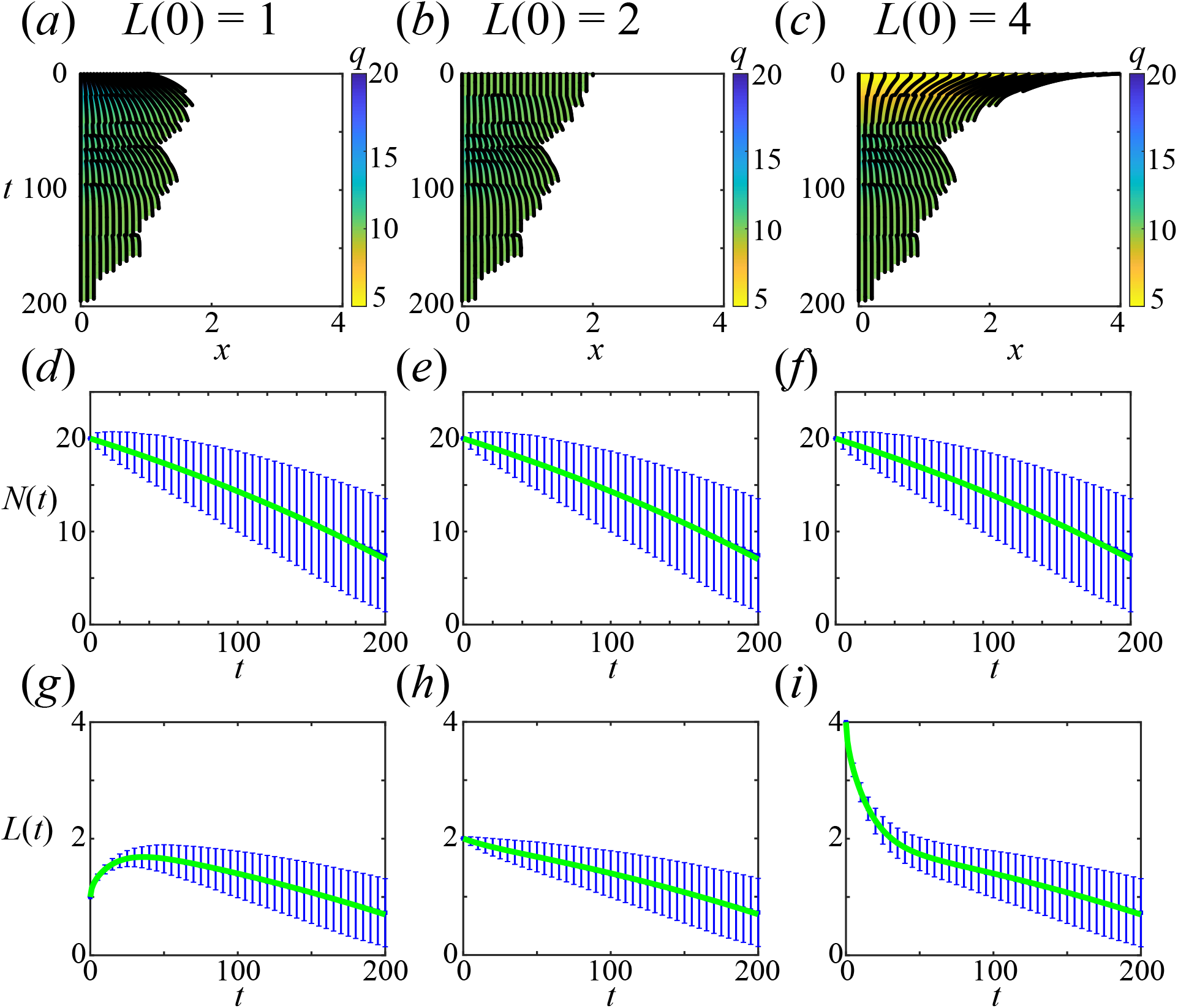
Cell-length-independent proliferation and chemically-independent cell detachment for *N* (0) = 20 and *k* = 10 with initial conditions: (a),(d),(g) compressed, (b),(e),(h) mechanical equilibrium, (c),(f),(i) stretched. (a)-(c) Kymographs with density, *q*(*x, t*), colouring. The average of 2000 discrete realisations (blue) are compared with the continuum model (green). (d)-(f) Evolution of total cell number, *N* (*t*). (g)-(i) Evolution of tissue length, *L*(*t*). Mechanical parameters: *k* = 10, *a* = 0.1, *η* = 1.

**Figure 13.**
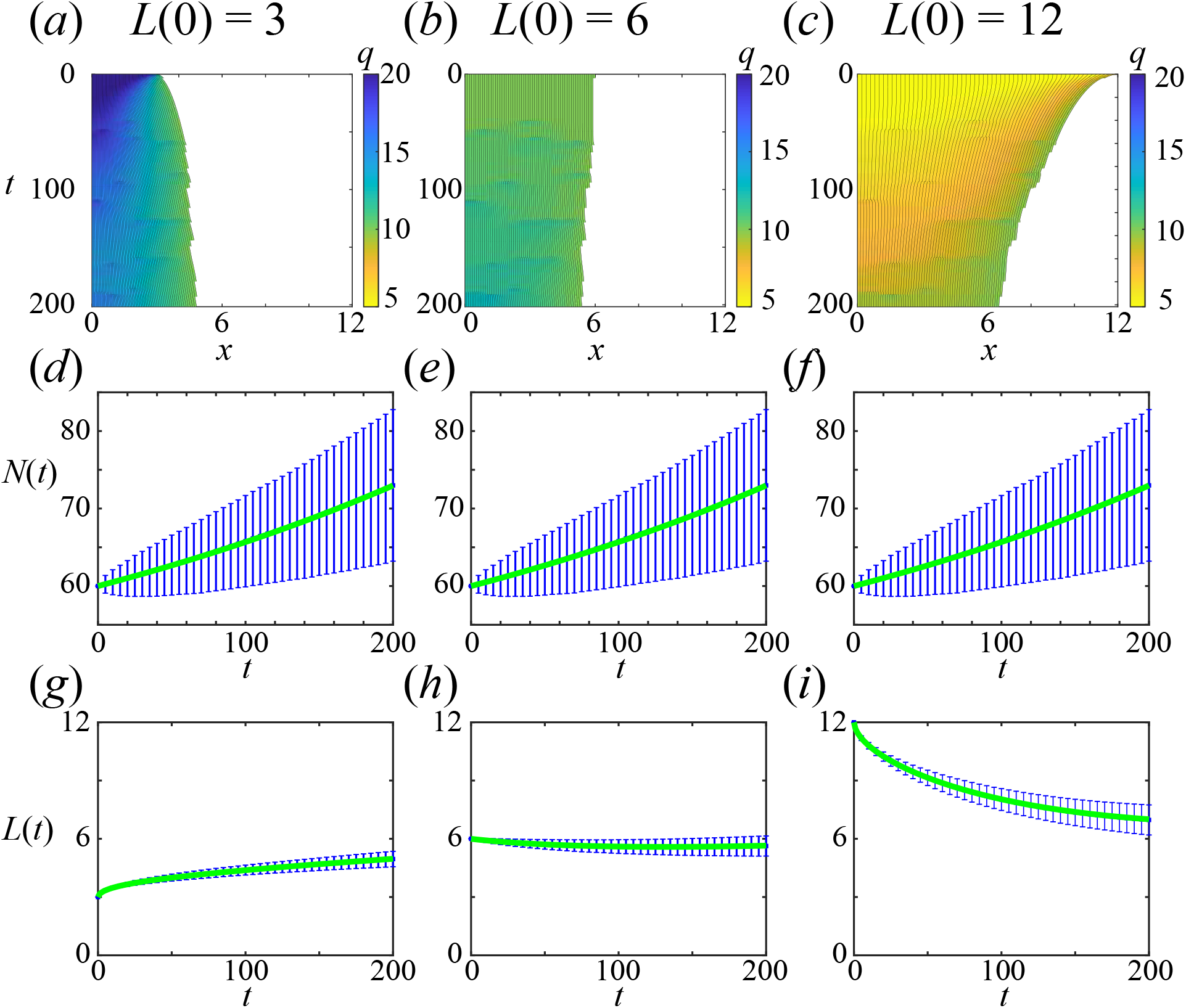
Cell-length-independent proliferation and chemically-independent cell detachment for N(0) = 60 and k = 10 with initial conditions: (a),(d),(g) compressed, (b),(e),(h) mechanical equilibrium, (c),(f),(i) stretched. (a)-(c) Kymographs with density, q(x; t), colouring. The average of 2000 discrete realisations (blue) are compared with the continuum model (green). (d)-(f) Evolution of total cell number, N(t). (g)-(i) Evolution of tissue length, L(t). Mechanical parameters: k = 10; a = 0:1; *η* = 1.

In Figures 11 and 14 we compare snapshots of the density, *q*(*x, t*), from the continuum model to the average of many discrete realisations and observe good agreement. Differences at the free boundary *x* = *L*(*t*) are a result of the length in each discrete realisation being different due to the stochastic proliferation and cell detachment mechanisms.

**Figure 14.**
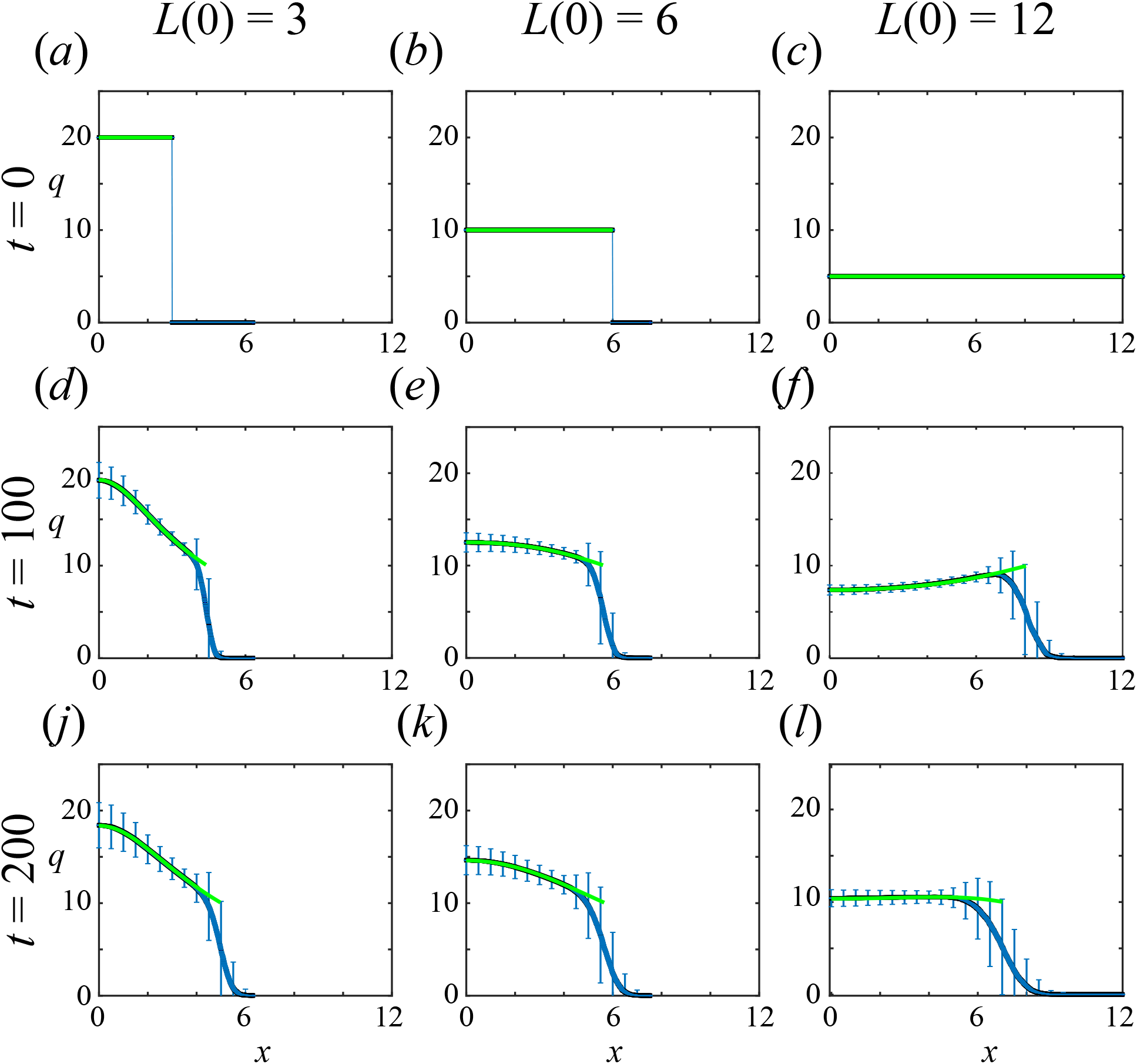
Density snapshots corresponding to Figure 13. The average of 2000 discrete realisations (blue) are compared with the continuum model (green) at times t = 0; 100; 200 for initial tissue lengths, L(0) = 3; 6; 12. Mechanical parameters: k = 10; a = 0:1; *η* = 1.

### D.3 Cell-length-dependent proliferation

Previously, with cell-length-independent proliferation starting with *N* (0) = 42 leads to unbounded growth in the continuum model, and mostly unbounded growth but sometimes extinction for realisations of the discrete model. Now we consider cell-length-dependent proliferation, where the initial tissue length can influence the long-term behaviour. Here we also assume chemically-independent cell detachment. Now with *N* (0) = 42, if the tissue is initially compressed then extinction is more likely (Figure 15a,c,e) whereas if the tissue is initially stretched then the tissue is more likely to eventually grow without bound (Figure 15(b),(d),(f)). Good agreement is observed between the average of many discrete realisations and the continuum description with differences due to being close to extinction (Figures 15(a),(c),(e)) and due to some discrete realisations crossing the critical length threshold while others do not (Figures 15(b),(d),(f)).

**Figure 15.**
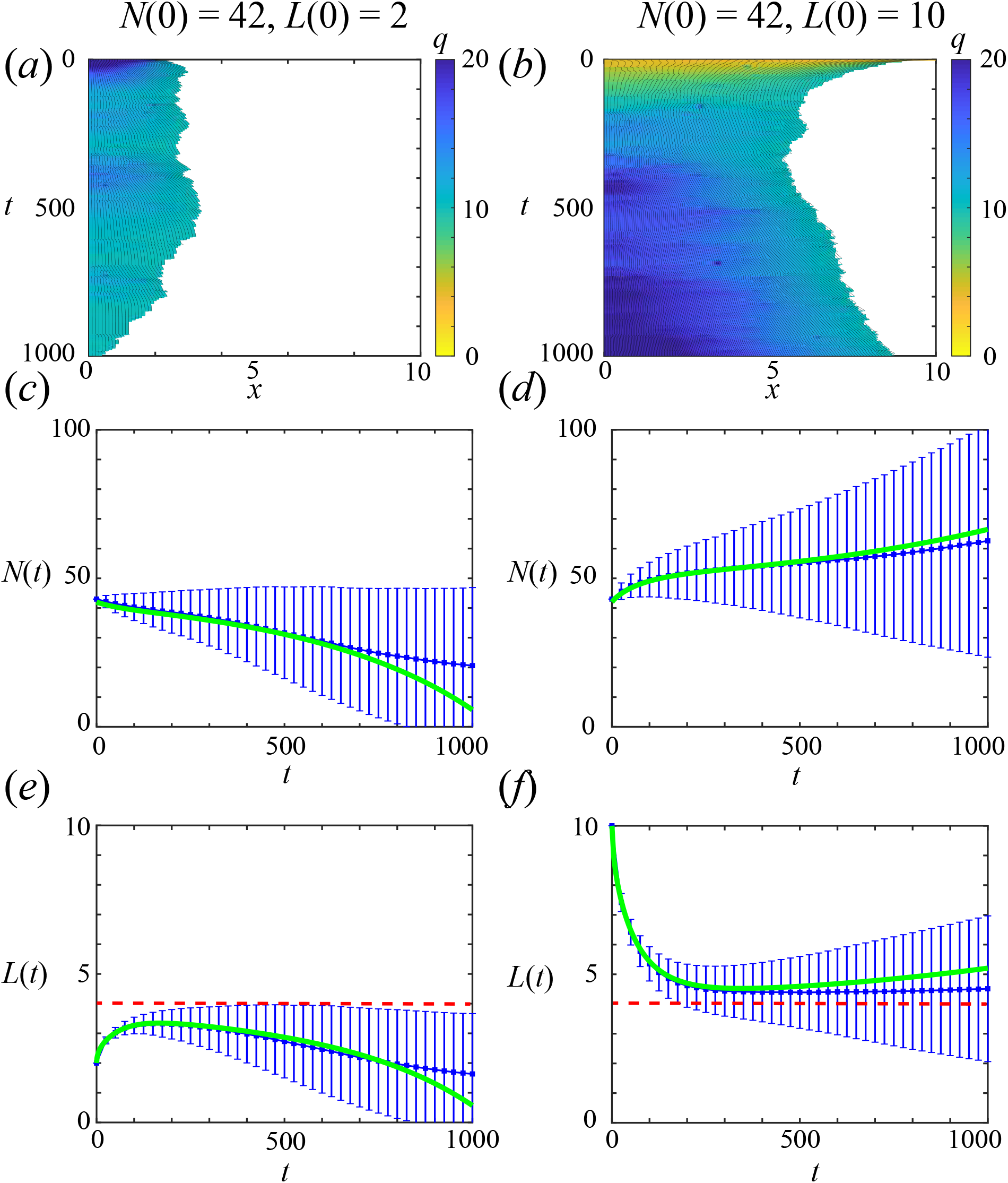
Chemically-independent cell detachment with linear cell-length-dependent proliferation mechanism. Two initial cell populations with *N* (0) = 42, the first uniformly compressed with *L*(0) = 2 and the second uniformly stretched with *L*(0) = 10 (a)-(b) Kymographs with density, *q*(*x, t*), colouring. The average of 2000 discrete realisations (blue) are compared with the continuum model (green). (c)-(d) Evolution of total cell number, *N* (*t*). (e)-(f) Evolution of tissue length, *L*(*t*). Red dashed line in (e)-(f) corresponds to the critical tissue length, *ω/β*. (e)-(f) Evolution of total cell number, *N* (*t*). Mechanical parameters: *k* = 1, *a* = 0.1, *η* = 1.

### D.4 Differences between the cell-based and continuum models: low *N* (*t*)

In the main manuscript, we state that the average of many cell-based realisations does not agree with the solution of the continuum model when *N* (*t*) is low and close to extinction. Here we provide further explanation.

In Figure 16(a), we extend the results from Figure 3 to *t* = 400. We observe that after approximately *t* = 200 there is a difference between the average of many discrete realisations and the continuum model. Similar behaviour is observed for the evolution of *L*(*t*) in Figure 16(b). These differences do not reduce when more simulations are performed. In Figure 16(c) we compare the solution of the continuum model with fifteen realisations of the discrete model and observe that many discrete realisations go extinct before the continuum model reaches *N* (*t*) = 0.

**Figure 16.**
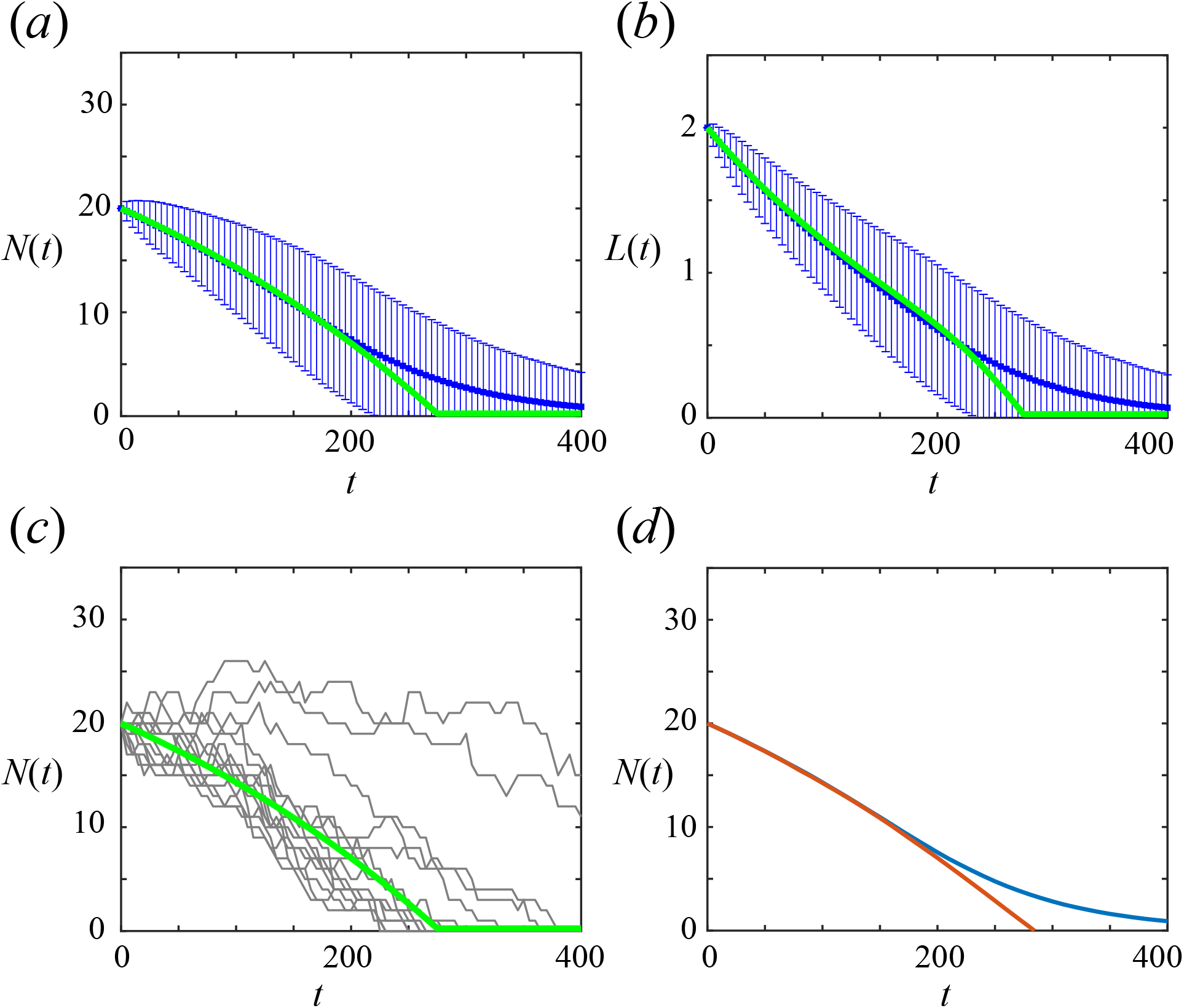
Differences near extinction between discrete and continuum results for total cell number and interface position. The average of 2000 discrete realisations (blue) are compared with the continuum model (green). (a) Evolution of total cell number, *N* (*t*). (b) Evolution of tissue length, *L*(*t*). (c) Evolution of number of cells, *N* (*t*), with 15 discrete realisations compared to solution from the continuum model. Mechanical parameters: *k* = 10, *a* = 0.1, *η* = 1. (d) Difference between discrete realisations stopped when *N* (*t*) = 0 (blue), which is physically realistic, and those where *N* (*t*) *<* 0 was allowed which is physically unrealistic (red).

In Figure 16(d) we simulate only the total cell number of the discrete model, which evolves stochastically according to *N* (*t* + d*t*) = *βN* (*t*) *− ω*, with two methods: i) whenever *N* (*t*) = 0 in an individual realisation it is stopped; ii) allow *N* (*t*) *<* 0. We find that if we allow *N* (*t*) *<* 0 then the discrete model matches results from the continuum model. However, allowing *N* (*t*) *<* 0 is physically unrealistic. Instead, stopping individual realisations when *N* (*t*) = 0 is physically realistic. Therefore, we must accept that the continuum model does not faithfully replicate the behaviour of the discrete model near extinction. Hence, the discrete model should be used when considering populations with low *N* (*t*).

### D.5 Chemically-dependent EMT: cell-length-independent proliferation

#### D.5.1 Small diffusivity, *D* = 10^*−*5^

In Figure 6(d),(g) of the manuscript we observe that there is a difference between the average of many discrete realisations and the continuum model. We explain that this is due to the well-mixed assumption for chemical concentration inside cells not being valid for small diffusivities. We show this in Figure 17. In Figure 17(a) we compare a snapshot of the chemical concentration from the continuum model and a discrete realisation at very early time, *t* = 0.15. In Figure 17(b) we compare the *c*(*L*(*t*), *t*) from the continuum model with *c*_*N*_ (*t*) from fifteen realisations of the discrete model, as these are used to calculate the rate of cell detachment. These results show that the continuum model reaches the concentration threshold much earlier than the discrete model which causes the number of cells in the continuum model to reduce faster than in the discrete model, hence explaining the difference in Figure 6(d),(g).

**Figure 17.**
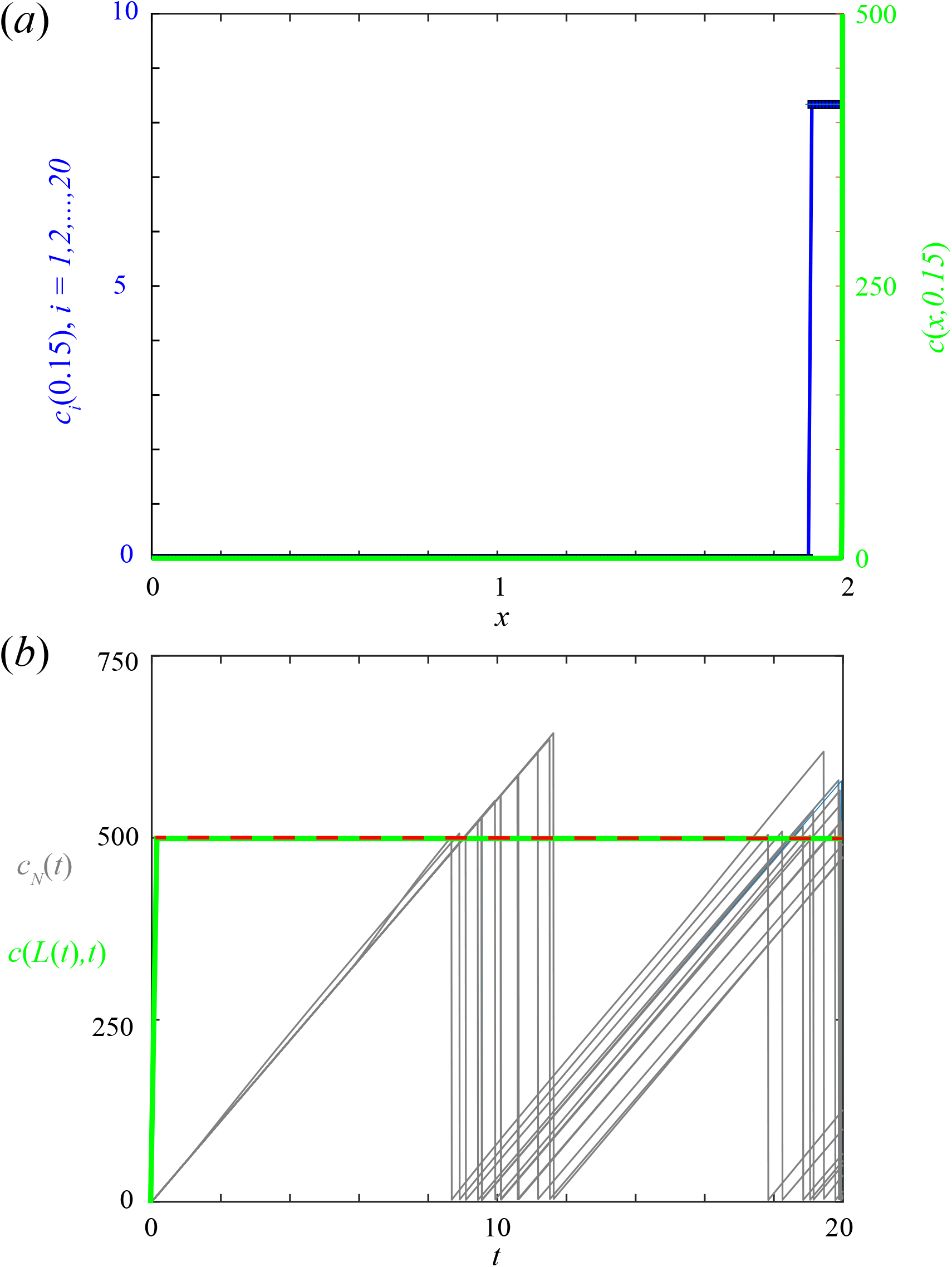
Continuum model (green) reaches concentration threshold (red-dashed) faster than individual realisations of the discrete model (grey). Chemically-dependent EMT with *D* = 10^*−*5^ and with cell-length-independent proliferation. (a) Concentration snapshot from the continuum model at time *t* = 0.15, *c*(*x*, 0.15) for 0 *< x < L*(1), compared to one realisation of the discrete model, where the chemical concentration in cell *i* is *c*_*i*_(0.15) for *i* = 1, 2, …, *N*. (b) Chemical concentration at the boundary node of the continuum model, *c*(*L*(*t*), *t*), compared to the chemical concentration of the boundary cell, *c*_*N*_ (*t*), for fifteen realisations of the discrete model. Here *N* (0) = 20. Mechanical parameters: *k* = 1, *a* = 0.1, *η* = 1

#### D.5.2 Starting closer to the chemical threshold with *D* = 1

In Figure 6(f),(i) of the manuscript we observe that there is a difference for *D* = 1. Here, we show that this difference is due to stochastic effects in realisations of the discrete model. Each discrete realisation has a different tissue length, *L*(*t*), resulting in the concentration threshold being reached at different times, which does not occur in the continuum model (Figure 18(a)).

**Figure 18.**
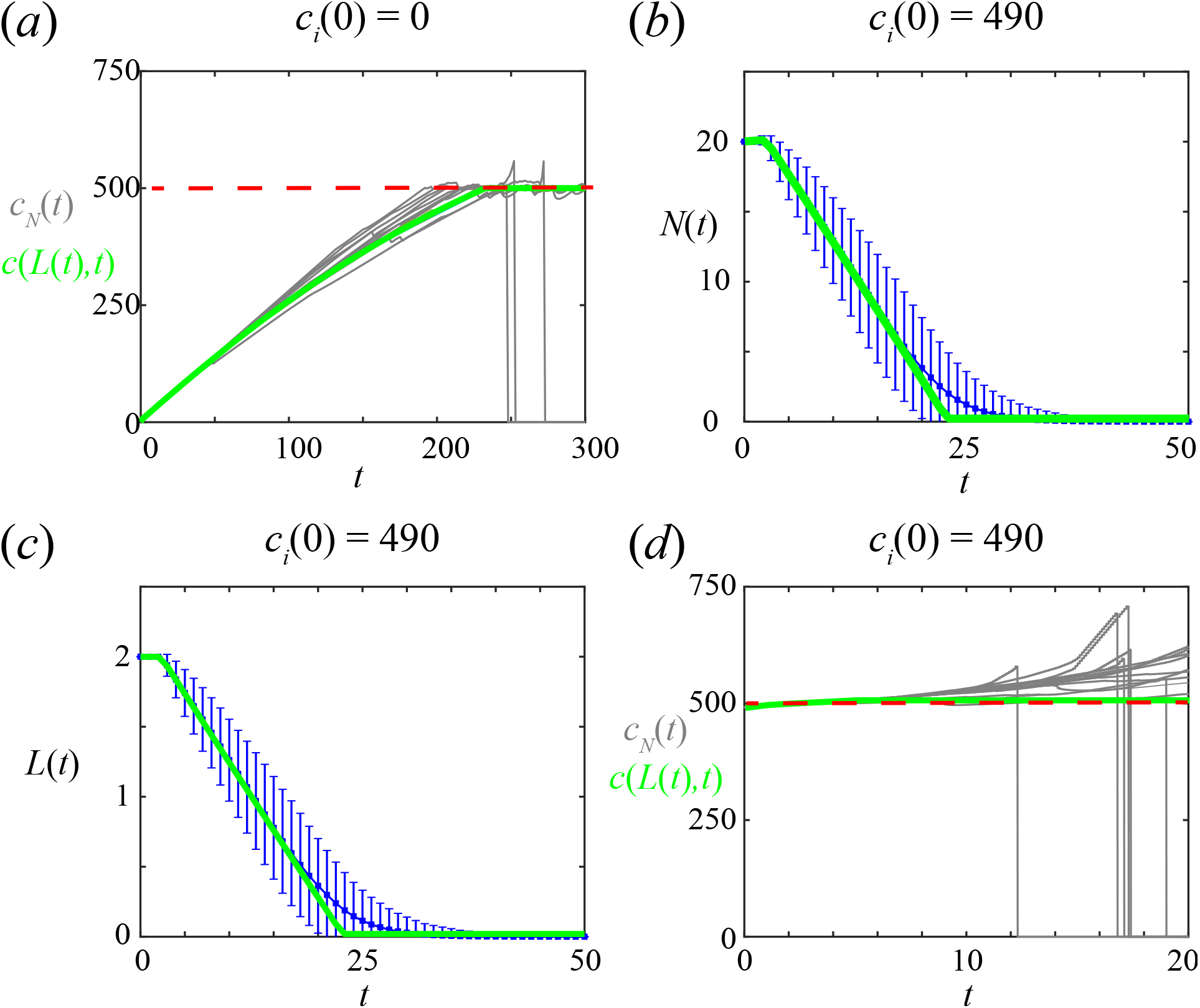
Comparison of results from continuum model and the average of many realisations of the discrete model for chemically-dependent EMT with *D* = 1 and cell-length-independent proliferation. (a) Evolution of the chemical concentration in the final cell, *c*_*N*_ (*t*), when *c*_*i*_(0) = 0 for *i* = 1, 2, …, 20. Some discrete realisations reach the concentration threshold, *C* = 500 (red-dashed), earlier than the continuum model. (b)-(c) The average of 2000 discrete realisations (blue) are compared with the continuum model (green). (b) Evolution of total cell number, *N* (*t*). (c) Evolution of tissue length, *L*(*t*). (d) Evolution of the chemical concentration in the final cell, *c*_*N*_ (*t*), when *c*_*i*_(0) = 490 for *i* = 1, 2, …, 20. Realisations of the discrete model reach the concentration threshold, *C* = 500 (red-dashed), at approximately the same time as the continuum model. Fifteen individual realisations of the discrete model are shown in (a) and (d). Mechanical parameters: *k* = 1, *a* = 0.1, *η* = 1.

If instead, we compare results when starting close to the chemical threshold, *C*, we find an improved match. Specifically, starting with *D* = 1 and *c*_*i*_ = 490 for *i* = 1, 2, …, 20, and *C* = 500, rather than with *c*_*i*_ = 0 for *i* = 1, 2, …, 20, we find an improved match (Figure 18(b)-(c)). We observe that different realisations of the discrete model reach the concentration threshold, *C* = 500, at approximately the same time as the continuum model (Figure 18(d)). This is due to the reduced time for stochastic effects in *N* (*t*) and *L*(*t*) to play a role.

### D.6 Sensitivity to *φ*

Chemically-dependent cell detachment is a two-step process: i) the boundary cell gains an invasive phenotype when the chemical concentration inside the boundary cell is above the chemical threshold, *C*; ii) the boundary cell detaches. We introduce a parameter *φ ∈* [0, 1] which defines the ratio of the average time in process i) as *φ/ω* and the average time in process ii) as (1 − *φ*)*/ω* (Figure 2). In the manuscript we present results for *φ* = 0.9 (Figure 6). Here, in Figure 19 we present results for *φ* = 0.1. As before, we choose parameters so that the average rate of cell detachment is the same as in previous models. To do this we keep the chemical threshold, *C*, fixed and vary the constant number of molecules per unit time supplied to the boundary cell from the external environment, *S*. Results for *φ* = 0.1 show improved agreement between the continuum model and the average of many discrete realisations in comparison to *φ* = 0.9. This is because the time to reach the chemical threshold is quicker and the time to stochastically detach is longer. Results for *D* = 10^−5^ (Figures 19(a),(d),(g)) and *D* = 10^−2^ (Figures 19(b),(e),(h)) look similar as they both reach the chemical threshold after a very short time. Note that when *φ* = 0 the chemically-dependent model is the same as the chemically-independent model.

**Figure 19.**
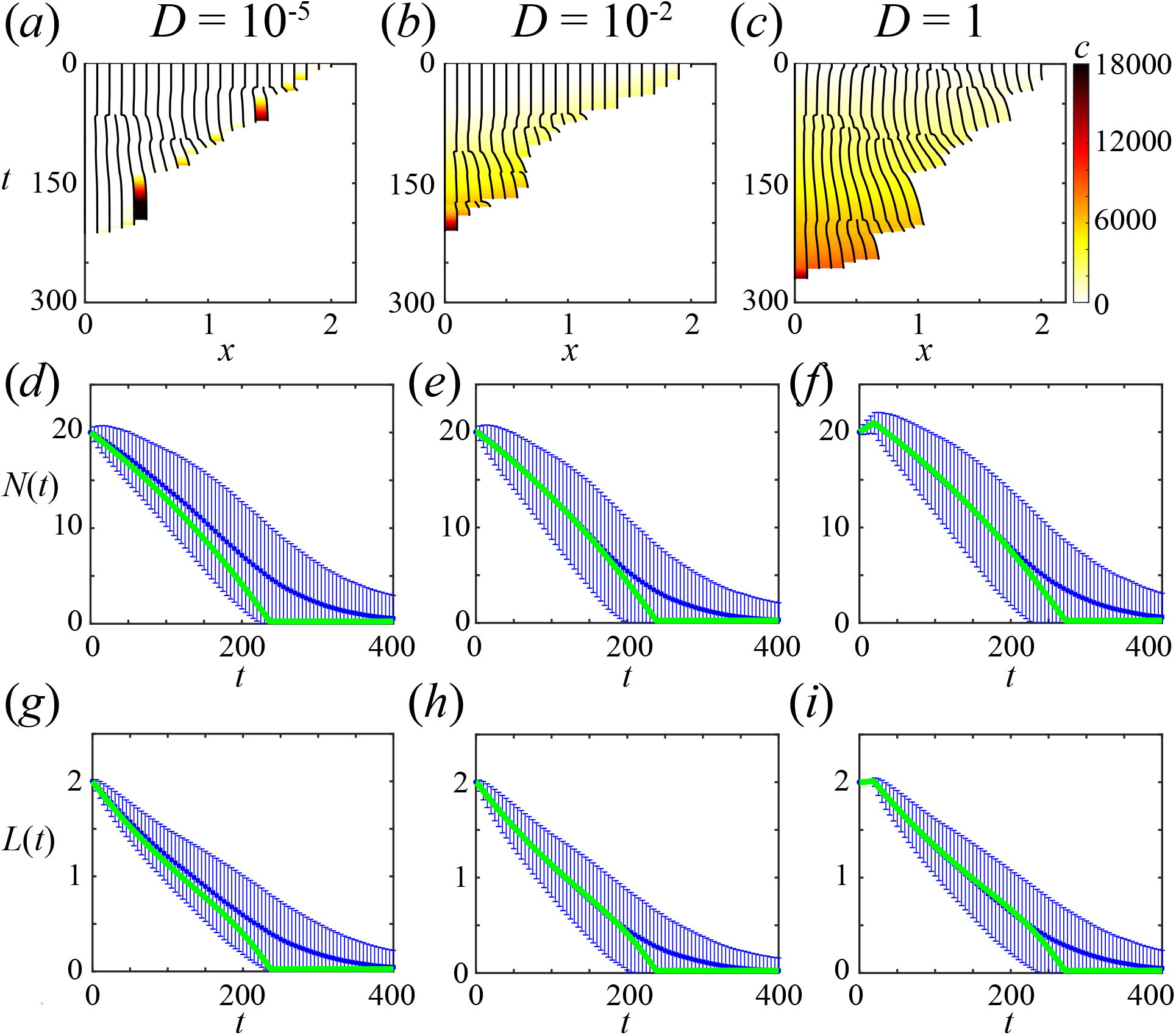
Sensitivity to *φ*. Figure 6, exploring diffusion delaying first EMT event, repeated with *φ* = 0.1. Cell detachment driven by chemically-dependent EMT with varying diffusivities and cell-length-independent proliferation mechanism. Cells initially at their resting cell lengths with initial cell populations *N* (0) = 20. Kymographs with chemical concentration, *c*(*x, t*), colouring shown for (a) *D* = 10^−5^, (b) *D* = 10^−2^, (c) *D* = 1. (d)-(i) The average of 2000 discrete realisations (blue) are compared with the continuum model (green). (d)-(f) Evolution of total cell number, *N* (*t*). (g)-(i) Evolution of tissue length, *L*(*t*). Mechanical parameters: *k* = 1, *a* = 0.1, *η* = 1.

In the above, we keep *C* fixed and vary *S*. Alternatively, one could vary *C* and keep *S* fixed, or vary both *C* and *S*. Furthermore, one could also keep both *C* and *S* fixed, and varying the rate of cell detachment. However, we do not show this here as then the average time for the boundary cell to detach is not the same as the average time to detach for previous models, which would not be a fair comparison.

### D.7 Chemically-dependent EMT: linear proliferation

In the manuscript we present results for cell detachment driven by chemically-dependent EMT and a cell-length-independent proliferation mechanism. Here we present results with a linear cell-length-dependent proliferation mechanism.

In Figure 20 we observe that when comparing an initially compressed tissue to an initially stretched tissue, with the same *N* (0), the time to reach the chemical threshold and the time for all cells to detach is shorter for the initially compressed tissue. This is due to proliferation being less likely in the compressed tissue with cell-length-dependent proliferation. This is also due to the chemical reaching the concentration threshold faster in the compressed tissue than in the initially stretched tissue, as there is less space for the chemical to diffuse in the compressed tissue.

**Figure 20.**
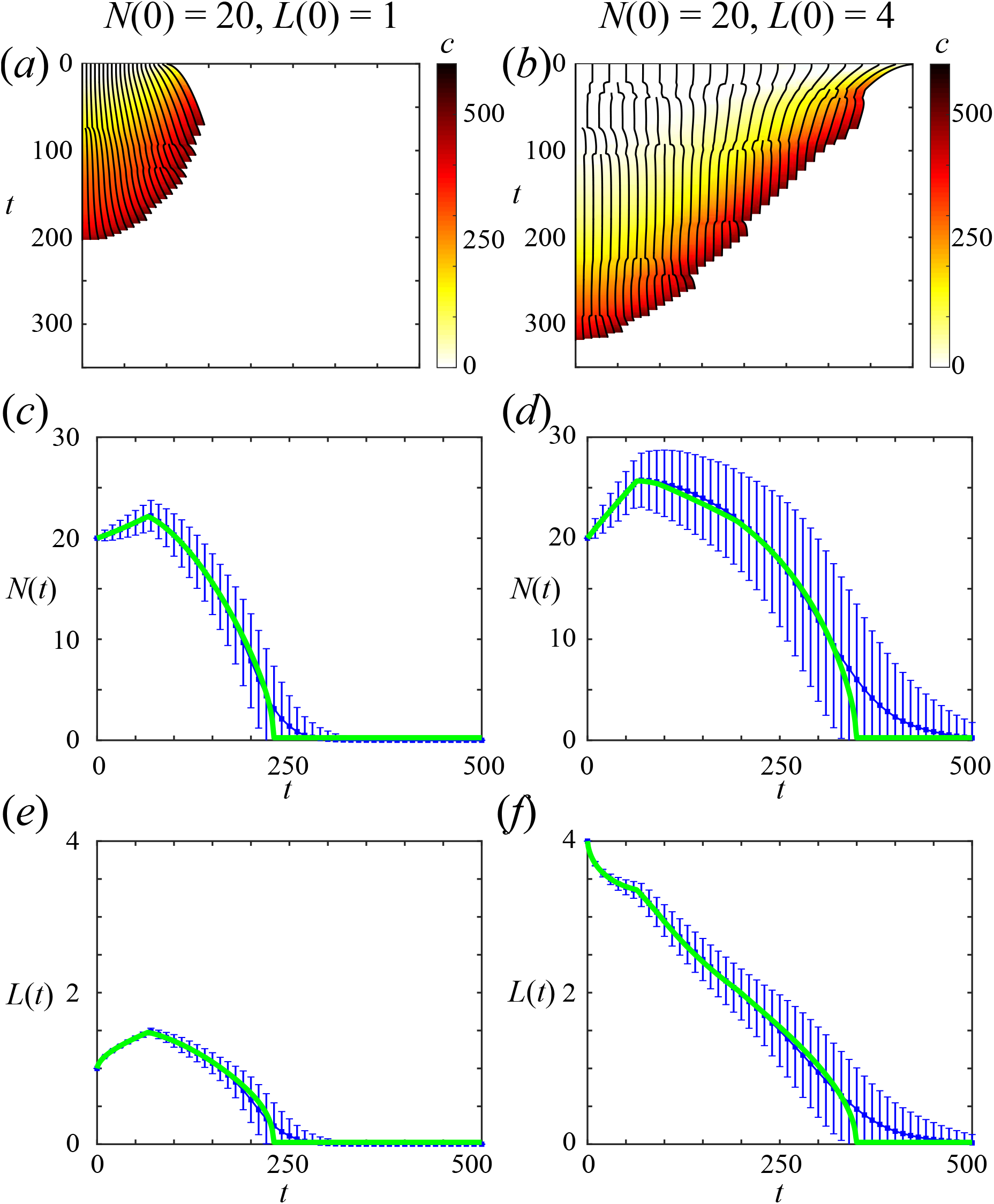
Cell detachment at driven by chemically-dependent EMT with *D* = 10^−2^ and with linear cell-length-dependent proliferation. Two initial cell populations with *N* (0) = 20, the first uniformly compressed with *L*(0) = 1 and the second uniformly stretched with *L*(0) = 4. (a)-(b) Kymographs with density, *q*(*x, t*), colouring. (c)-(f) The average of 2000 discrete realisations (blue) are compared with the continuum model (green). (c)-(d) Evolution of total cell number, *N* (*t*). (e)-(f) Evolution of tissue length, *L*(*t*). Mechanical parameters: *k* = 1, *a* = 0.1, *η* = 1.

## E Diffusive equilibrium at all times

If the diffusivity of the EMT-inducing chemical is very high it is reasonable to assume that the chemical in the tissue is at diffusive equilibrium at all times. This simplifies the analysis as every cell in the tissue experiences the same concentration at time *t*, which we denote *c*(*t*). This can be useful to understand possible long-term behaviours.

To proceed, we make a further assumption that cells are always at mechanical equilibrium and consider the continuum model. As cells are at mechanical equilibrium, the cell-length-independent and cell-length-dependent proliferation mechanisms are equivalent, so before any cell detachment events occur *N* (*t*) = *N* (0) exp(*βt*), *L*(*t*) = *N* (*t*)*a*, and *c*(*t*) = *St/L*(*t*). Then for cell detachment to occur at least once we require that *c*(*t*) *> C* which is equivalent to requiring *N* (0) *< S/* (*Caβ* exp (1)) = *N*_*I*_.

So if *N* (0) *> N*_*I*_ the tissue grows without bound and there is no EMT and no cell detachment, and *N* (*t*) evolves according to d*N* (*t*)*/*d*t* = *βN* (*t*). However, if *N* (0) *< N*_*I*_ the time to reach the concentration threshold, *t*_*C*_, is the solution of *t*_*C*_*/* exp(*βt*_*C*_) = *CN* (0)*a/S*. While *c*(*t*) *> C, N* (*t*) evolves according to d*N* (*t*)*/*d*t* = *βN* (*t*) − *φω* and if the concentration decreases below *C* then *N* (*t*) evolves according to d*N* (*t*)*/*d*t* = *βN* (*t*). This results in four possible behaviours:

1. unbounded tissue growth without EMT and cell detachment (Figure 21(a),(b));
2. unbounded tissue growth with some EMT (Figure 21(c),(d));
3. eventual tissue homeostasis and constant EMT (Figure 21(e),(f));
4. eventual tissue extinction due to EMT (Figure 21(g),(h)).

**Figure 21.**
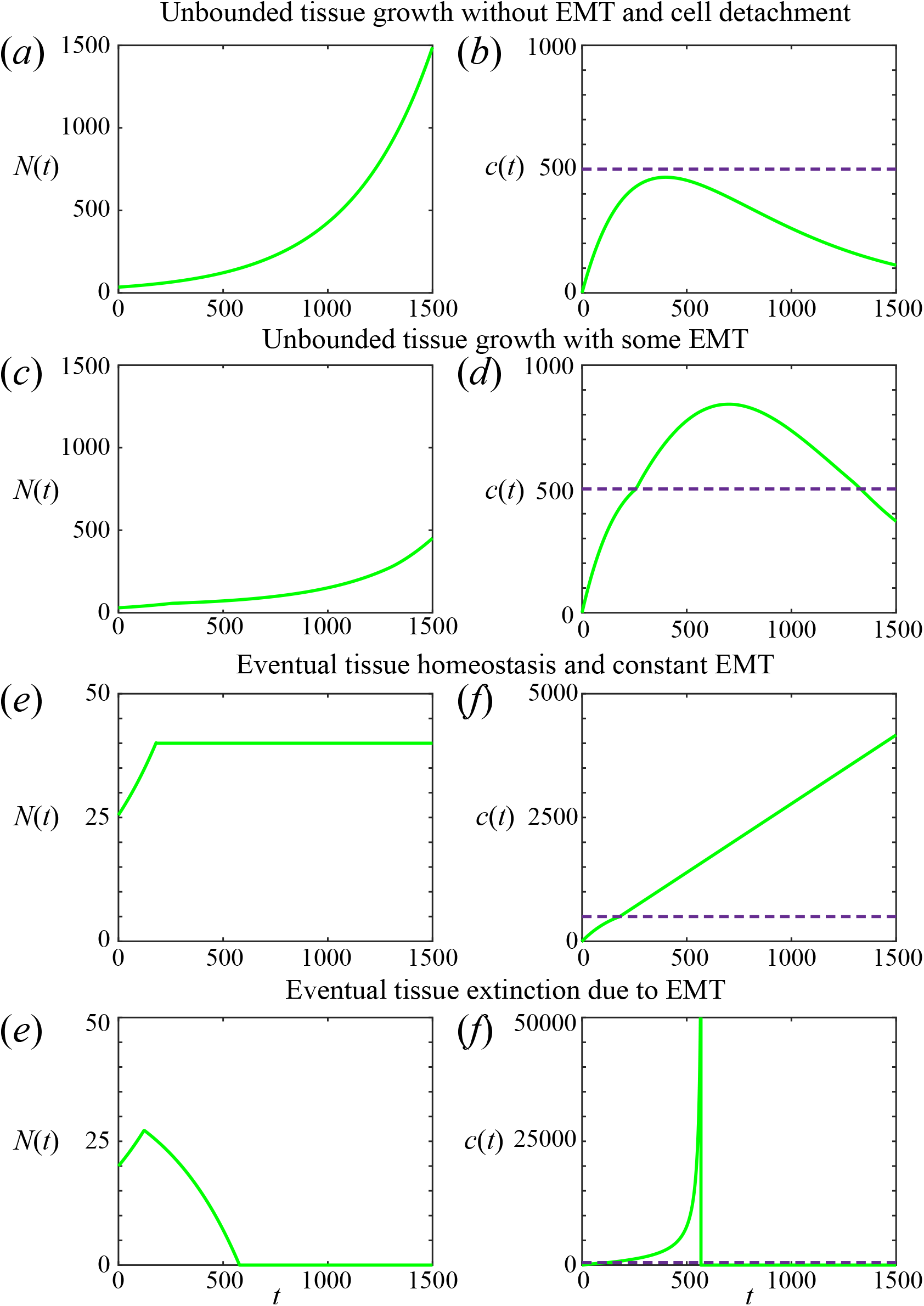
For diffusive equilibrium at all times and instantaneous mechanical relaxation there are four possible behaviours. (a) Unbounded tissue growth without EMT and cell detachment. (b) Unbounded tissue growth with some EMT. (c) Eventual tissue homeostasis and constant EMT. (d) Eventual tissue extinction due to EMT. Purple dashed line corresponds to chemical threshold *C*. Results shown for *S* = 50*/*9, *C* = 500, *φ* = 0.9, *k* = 1, *a* = 0.05, *β* = 0.0025, *ω* = 0.1.

## Notes

https://github.com/ryanmurphy42/Murphy2020b

### Competing Interest Statement

The authors have declared no competing interest.

